# Necrosignaling: Cell death triggers antibiotic survival pathways in bacterial swarms

**DOI:** 10.1101/2020.02.26.966986

**Authors:** Souvik Bhattacharyya, David M. Walker, Rasika M. Harshey

## Abstract

Swarming is form of collective bacterial motion enabled by flagella on the surface of semi-solid media^1^. Many bacterial species exhibit non-genetic adaptive resistance to a broad range of antibiotics only when swarming (SR)^2-4^. While the swarming population as a whole survives antibiotic challenge, it nonetheless sustains considerable cell death^5^. In this study focused mainly on *E. coli*, an initial analysis of antibiotic-induced killing patterns of swarm vs planktonic cells indicated that death of a sub-population is beneficial to the swarm in promoting SR. Introduction of pre-killed cells into a swarm indeed enhanced SR, allowing us to purify the SR factor from killed cells, and to establish that cell death is directly involved in SR. The SR-enhancing factor was identified to be AcrA, a periplasmic component of a tripartite RND efflux pump^6^, of which the outer membrane component TolC is shared by multiple drug efflux pumps^7^. We show that AcrA stimulates drug efflux in live cells by interacting with TolC from the outside. This stimulus acts as a signal to activate efflux in the short term, and to induce the expression of other classes of efflux pumps in the long term, amplifying the response and establishing SR. We call this phenomenon ‘necrosignaling’, and show the existence of species-specific necrosignaling in both Gram-positive and Gram-negative bacteria. Our results have implications for the adaptive resistance of other surface-associated bacterial collectives such as biofilms. Given that such resistance is known to be an incubator for evolving genetic resistance^8^, our findings may also be relevant to chemotherapy-resistant cancers.

## Main

Bacteria employ many appendages for movement and dispersal in their ecological niches^9^. Flagella-enabled swarming offers bacteria a competitive advantage in both ecological and clinical settings^10,11^. An unexpected and clinically relevant property of swarms is their SR at conditions lethal to free-swimming planktonic cells of the same species^2-5,12^. SR depends on high cell densities associated with swarms^5^, and is phenomenologically similar to adaptive resistance of bacterial biofilms^13^. While several terminologies such as resistance, tolerance and persistence have been used to describe the resistance phenotype observed under various settings^8,14^, a shared attribute of a majority these phenotypes is their association with reduced metabolism and slow bacterial growth^15^. In stark contrast, bacterial swarms are metabolically active and grow robustly. We will therefore employ a new term STRIVE (*<u>s</u>*warming *w*ith *<u>t</u>*emporary *<u>r</u>*esistance *<u>i</u>*n *<u>v</u>*arious *<u>e</u>*nvironments; SR for brevity) to distinguish swarming-specific from slow growth-related adaptive resistance.

## Death of a Sub-population May be Linked to STRIVE

While swarming across media with antibiotics, a substantial fraction of dead cells were observed in several bacterial species^5^. A potential role for the dead cells in enabling SR by forming a protective barrier around the live cells was ruled out^16^; instead, live cells were seen to migrate away from the antibiotic source, leading to a suggestion that cell death emanating around this source releases an avoidance ‘signal’ that contributes to SR^16^. Programmed bacterial cell death, where death of a subpopulation benefits the community by providing nutrients, has been reported by several studies^17,18,19^. The aim of present study, however, was to test if cell death within a community generates a distress signal to induce SR pathways for survival. Initially, we monitored the antibiotic susceptibility of planktonic vs swarm cells of *E. coli*. Exposure to a broad concentration range of the antibiotic kanamycin (Kan) produced distinct survival patterns (Fig. S1a, b). At low concentrations (e.g, Kan^2.5^; 2.5μg/ml), and during the early phase of growth (0.5 h), swarm cells were surprisingly more susceptible to killing than planktonic cells, a trend maintained for up to Kan^10^ (Fig. S1a). This trend reversed at both higher Kan concentrations as well as later times, where planktonic cells became more susceptible. We hypothesized that the higher initial susceptibility of swarm cells could be indicative of heterogeneity in the population, with a sub-population that was more sensitive. This rationale was tested *in silico* by simulating the survival patterns of each sub-population as a first order decay where the survival ‘rate’ is analogous to that population’s susceptibility to the antibiotic (Fig. S1a and SI Text). A homogeneous cell population with a singular response rate reasonably modeled the planktonic data (MSE range: 10^−6^-10^−3^), but failed to model the swarm data (MSE range: 10^−3^-10^−2^) (Fig. 1a). Conversely, when a heterogeneous population composed of two sub-populations differing in initial rates of death were introduced into the simulations, the resultant curves successfully fit the experimental data as determined by MSE scores 10 to 1000 times lower (MSE range: 10^−7^-10^−5^) compared to the homogeneous population. The fits between the two population models for both swarming and planktonic cells were determined to be statistically different by the Wilcoxon rank sum test (Fig. 1a). Taken together, the data indicate the existence of two distinct sub-populations with the swarm, one more susceptible to the antibiotic. Given that the experimental MIC for Kan for swarm cells was almost double that of planktonic cells (Fig. S1c), we hypothesized that the more susceptible population might serve as an early distress signal to elicit SR in the community.

**Figure 1.**
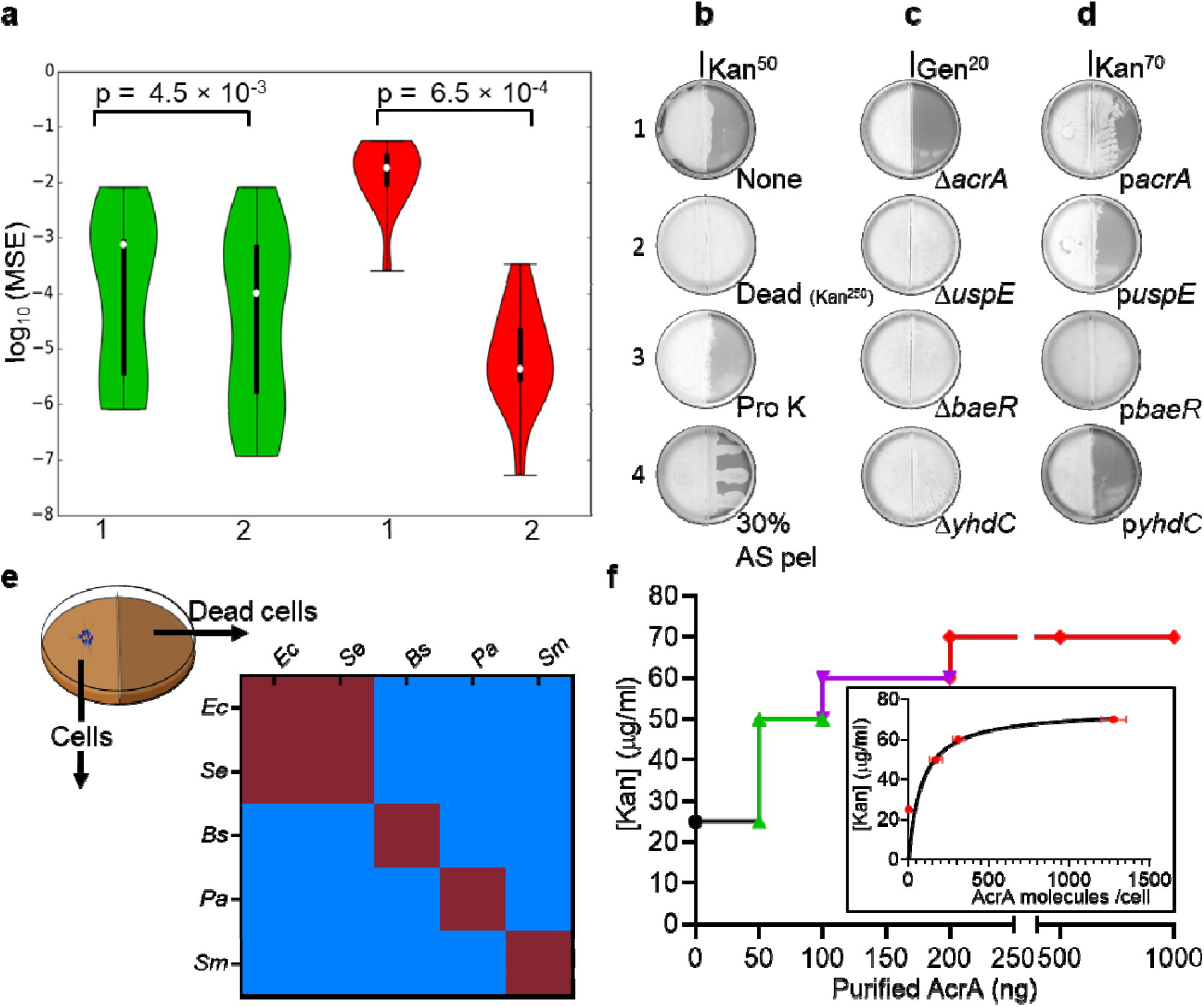
AcrA released from dead bacteria acts as a ‘necrosignal’ to promote STRIVE in *E. coli*. **a**. Grap showing distribution of the mean squared errors (MSE) between simulated (dotted lines) and experimental (solid lines) killing curves shown in S1A of planktonic (green) and swarm (red) cells; labels 1 and 2 indicate homogeneous and heterogeneous populations respectively. The p-values indicated were calculated from a Wilcoxon rank sum test between the homogeneous and heterogeneous populations in planktonic and swarm cells. **b-d**. Border-crossing assays that established the identity of the necrosignal. Wild-type *E. coli* were inoculated in the left chamber in every case, whereas material applied to the right chamber is indicated below each plate. **b**. None, no cells applied; Dead (Kan^250^), dead cells killed by Kan^250^; Pro K, dead cell supernatant treated with Proteinase K; AS pel, pellet fraction after treating cell supernatant with ammonium sulfate. Kan, Kanamycin; Gen, Gentamycin. **c**. Gene deletions (Δ). All gene deletions were made with a Kan cassette, so the cells were pre-killed with Gentamycin (Gen^50^), and tested for swarming on Gen^20^. **d**. Gene overexpression from ASKA library plasmids (p). These strains were pre-killed with Kan^250^. **e**. Chart showing the species specificity of necrosignaling. *Ec, Escherichia coli*; *Se, Salmonella enterica*; *Bs, Bacillus subtilis*; *Pa, Pseudomonas aeruginosa*; *Sm, Serratia marcescens*. Columns: bacterial species inoculated in the left chamber. Rows: bacterial species providing the dead cells applied on the right chamber, Maroon, SR+ response; Blue, SR-response. **f**. Swarming response to indicated Kan concentrations with increasing AcrA applied to the right chamber. The response was saturated at Kan^70^. Inset: minimum number of AcrA molecules estimated (from the protein concentration) to be required for STRIVE (estimated from CFUs) at the indicated Kan concentrations. The data (red dots) were fit to an exponential function (black line).

## Dead Cells Release AcrA as a Necrosignal

To directly test whether cell death acts as a signal, the border-crossing assay was primarily employed^5^ as diagrammed in Figure S2a, where in a divided petri plate containing swarm media, only the right chamber has the antibiotic. Wild-type *E. coli* inoculated on the left, swarms over the right chamber with Kan^25^ (not shown) but not with Kan^50^ (Fig. 1, b1). When *E. coli* cells killed by Kan^250^ were applied to the right chamber, the wild-type population could now colonize Kan^50^ (Fig. 1, b2). Although cells killed with Kan promoted migration over Kan^50^, the enhanced SR response was independent of the killing method, with the exception of heat (Fig. S3, a-b). The response to killed cells was sustained, in that the swarm retained its ability to STRIVE even after exiting a zone of dead cells (Fig. S4).

The heat-sensitive nature of the SR-factor (Fig. S3b) suggested that it might be isolatable. To this end, cell extracts prepared from Kan^250^-treated cells were assayed and showed activity in the supernatant fraction (Fig. S3b). The activity was resistant to DNaseI and RNaseI (Fig. S3c), but sensitive to protease (Fig. 1, b3). A 30% ammonium sulfate precipitate, when resuspended in buffer and applied as lines, promoted the swarm to track along these lines (Fig. 1, b4). We will henceforth refer to this active factor as the ‘necrosignal’, and its ability to promote SR as ‘necrosignaling’. We found necrosignaling to be operative in other bacterial species as well (Fig. 1e). However, except for *E. coli* and *Salmonella*, where killed cells of one species promoted a reciprocal response in the other (Fig. 1e, maroon areas), species-specificity of the response was evident in *Bacillus subtilis, Pseudomonas aeruginosa*, and *Serratia marcescens* (Fig. 1e, blue areas). Given that *E. coli* and *Salmonella* have an interchangeable response, we used both bacteria to purify and determine the common identity of the necrosignal (Fig. S5). MS/MS analysis of the active fractions obtained after the final purification step yielded 5 common proteins (Fig. S6; AcrA, UspE, BaeR, YdhC, Crp). All subsequent experiments were performed with *E. coli*.

Deletion and overexpression analysis helped narrow down the necrosignal candidate (Fig. 1c,d). In both analyses, candidate strains were pre-killed and applied to the right chamber. Only Δ*acrA* abolished the SR response to Kan^50^ (Fig. 1c; data for *crp* not shown). Conversely, when overexpressed, both AcrA (p*acrA*) and BaeR (p*baeR*) promoted SR to Kan^70^ (Fig. 1d). BaeR is a positive regulator of *acrAD* operon^20^. Given that Δ*baeR* did not abolish the response, AcrA is most likely the necrosignal. Purified AcrA (flanked by His- and FLAG-epitope tags), showed a concentration-dependent SR response, plateauing at Kan^70^ (Fig. 1f). The estimated number of AcrA molecules per cell required to elicit a response increased exponentially with increasing antibiotic (Fig. 1f, inset) reaching a plateau, suggesting specificity (non-specific binding is generally linear^21^). When purified AcrA was added to a planktonic culture treated with Kan near its MIC_99_, however, only a modest increase in survival was observed, unless cell density was increased (Fig. S7), but even then, swarm cells had a better response, indicating that high cell density is important but not sufficient to account for STRIVE, and that the physiological state of the swarm might be an important contributor as well.

### Mechanism of AcrA Necrosignaling

AcrA is the periplasmic component of the RND efflux pump AcrAB-TolC (and AcrAD-TolC)^22^. To elucidate whether the activity of this surprising candidate for the necrosignal was independent of outer membrane (OM) protein TolC, we tested the effect both deleting and overexpressing *tolC*. Absence of TolC in the active swarm annuled the SR response to AcrA applied on the right (Fig. 2, a1), while overexpression of TolC restored it (Fig. 2, a2). However, absence of TolC in killed cells applied on the right did not affect the response outcome (Fig. 2, a3). Taken together, these results indicate the necrosignaling activity of AcrA is dependent on the presence of TolC in the live cells, and suggest that AcrA released from dead cells may bind to TolC from the outside. We next sought to identify residues critical for the response in both AcrA and TolC. For TolC, we took advantage of reports that TolC serves as a receptor for certain colicins^23^ and bacteriophages^24^, using residues important for phage binding as a guide for our mutagenesis. As shown in Figure 2b and 2c (see Fig. S8 for primary data), a subset of the residues tested were also essential for the SR response. For AcrA, deletion of 72 and 75 residues from its C- and N-terminus respectively, did not affect its activity, but perturbation of its helix-turn-helix (HTH) motif or nearby residues eliminated it, suggesting that the HTH region of AcrA is critical. To test binding of AcrA to TolC exposed on the OM, we marked AcrA with Qdot^705^-labeled anti-FLAG antibody, and labeled the OM with FM-143. Microscopy images show localization of AcrA to the OM (Figs. 2d and S9a). Deletion and overexpression of *tolC* led to loss and restoration of AcrA binding to the OM, respectively (Figs. 2e and S9b). Critical TolC residues as deduced from plate assays (Fig. 2c) were verified in the binding assay by the observation that *tolC*^S257A^ but not in *tolC*^R55A^ supported AcrA binding (Figs. 2e and S9b). The external localization of AcrA was confirmed by trypsin digestion (Figs. 2e and S9c). Taken together, both genetics and microscopy confirm that AcrA binds TolC externally to elicit necrosignaling.

**Figure 2.**
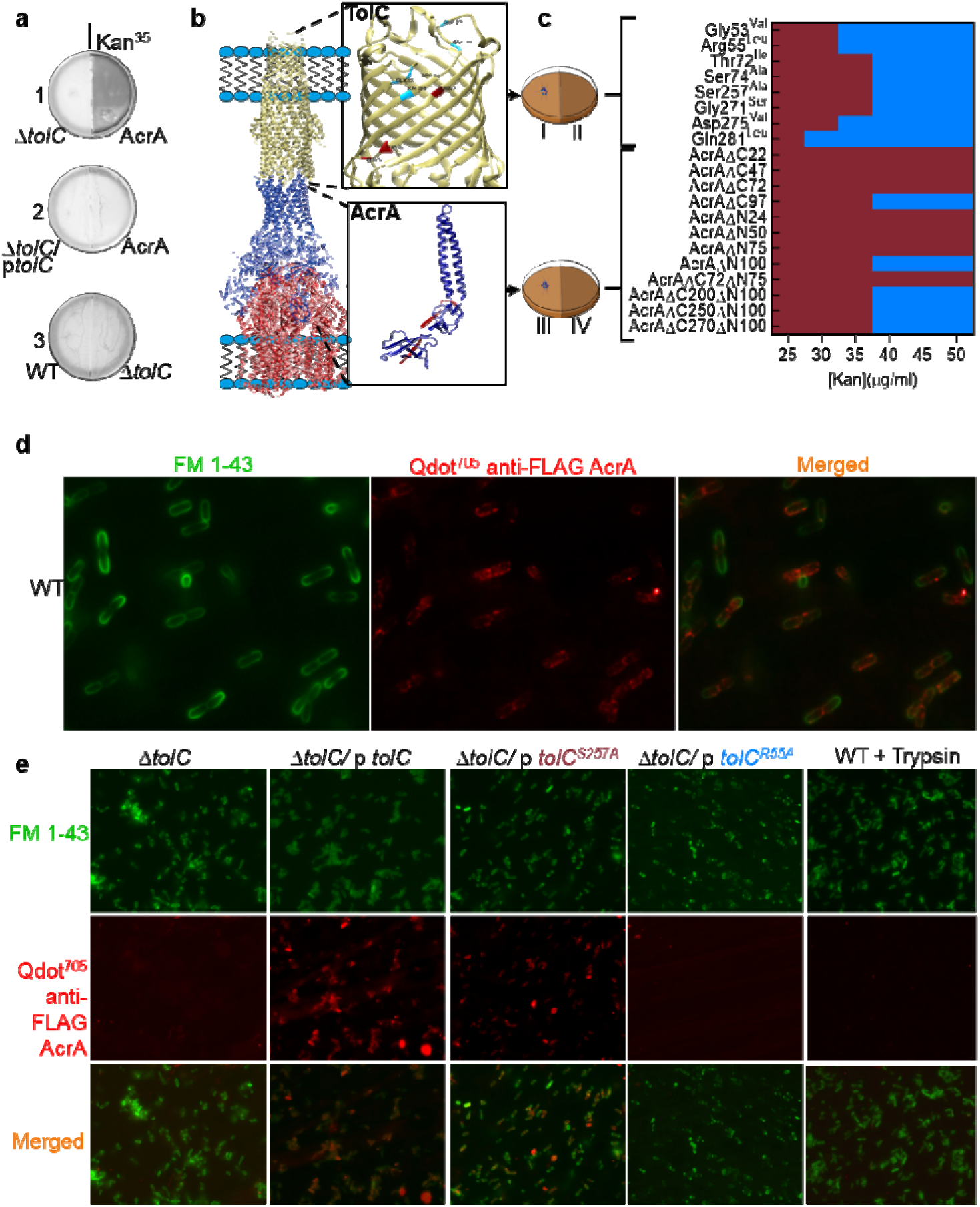
AcrA, a periplasmic component of RND drug efflux pumps, binds TolC externally as a necrosignal. **a**. The AcrA response requires TolC. Genotypes of *E. coli* strains inoculated on the left or proteins/pre-killed cells applied on the right are indicated below the plates (p, plasmid). The *tolC* deletion reduces SR to Kan^25^, hence the use of Kan^35^ in these plates **b**. A model of the AcrAB-TolC efflux pump (PDB ID: 5NG5) visualized and drawn (not to scale) using Chimera, with TolC and AcrA components enlarged. The enlarged view for TolC shows the residues mutated in this study, with maroon and blue indicating SR+ and - outcomes, and similar colors in AcrA indicate the importance of the HTH region for SR+ activity. **c**. Summary of data delineating residues in TolC, and regions in AcrA, important for STRIVE (see Fig. S8 for primary data). For TolC analysis, *tolC* mutants expressed from plasmids in a Δ*tolC* strain were inoculated on the left (I), with purified AcrA on the right (II). For AcrA, WT cells were inoculated on the left (III), and pre-killed cells expressing AcrA deleted for its indicated C-terminal or N-terminal residues were applied on right (IV). SR response color scheme as in Fig. 1e. **d**. Imaging of AcrA binding to the outer membrane of WT swarm cells. QDot^705^-labed anti-FLAG antibody was used to label AcrA-FLAG, and FM-143 dye to label the outer membrane, as described in Methods. The entire field of view is shown in Fig. S9a. **e**. AcrA-TolC co-localization in representative SR+ and SR-TolC mutants identified in c and treated as in d. See Fig. S9b for brightfield images. The last panel is a control to show that AcrA is not internalized. Here, WT swarm cells were pre-incubated with AcrA and subsequently treated with trypsin followed by addition of the Qdot (see Methods). See Fig. S9c for western blot analysis of these samples.

### Mechanism of STRIVE

Bacterial swarms alter their gene expression patterns and physiology compared to their planktonic counterparts^11,25,26^. To determine if necrosignaling impacts gene expression, RNA-Seq analysis was performed. Expression of 566 genes, many of which ontologically classified to motility, membrane function, and energy metabolism, showed specific changes in swarm cells (Figs. 3a and S10). Efflux pumps and transport functions were upregulated, with further increases in expression upon addition of antibiotics and of AcrA. Expression of OM porins genes were down in the swarm and those of ROS catabolism genes were up. These patterns suggest that swarms are pre-programmed to be less permeable^27^, more prepared to efflux, and to tolerate antibiotic stress by upregulating ROS pathways^28,29^. These inferences were verified in several ways. The Nile Red assay for efflux^30,31^ confirmed a higher rate of efflux, stimulated by AcrA (Fig. 3b). The Alamar Blue assay for membrane permeability^30,32^ showed lower permeability, unaffected by AcrA (Fig. 3c). Next, we directly measured the intracellular concentration of antibiotics using two methods – spectrometry employing the fluorescent antibiotic Fleroxacin^33^, and disc diffusion assay for the thermostable Kanamycin^34^. The concentration of both antibiotics was significantly lower in swarm cells compared to planktonic cells (Figs. 3d and S11), consistent with the data in Figure 3b and 3c. The RNA-Seq data was further validated by genetics. Overexpression of selected genes involved in ROS catabolism, iron transport (important for the regeneration Fe-S clusters of ROS catabolizing enzymes^35^ and during swarming^36^), and efflux pumps, increased SR, whereas overexpression of porins decreased it (Fig. S12a). Deletions of many efflux pump components and their regulators also decreased SR (Fig. S12b).

**Figure 3.**
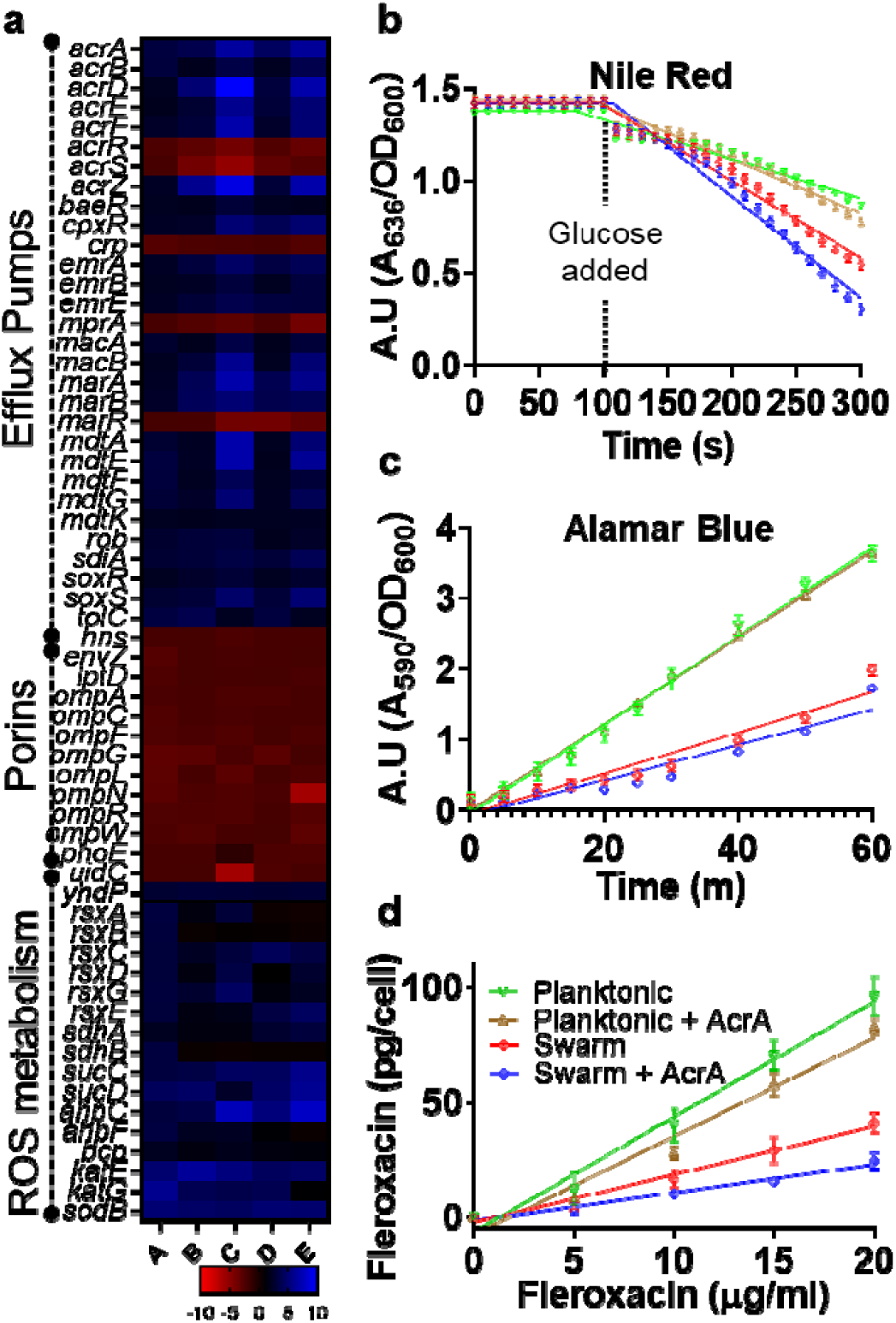
Mechanism of STRIVE. **a**. Comparison of log_2_ fold changes in gene expression. A) Swarm vs Planktonic, B) Swarm+Kan^20^ vs Swarm, C) Swarm+Kan^20^+AcrA vs Swarm, D) Swarm+Cip^2.5^ vs Swarm, E) Swarm+Cip^2.5^+AcrA vs Swarm. The concentration of AcrA was 0.1 μg/ml, and the log_2_ fold change cutoff value was 2. About 25 genes encoding efflux pump components representing all five classes of pumps and their regulators were upregulated in swarm cells (∼ 2 - 3 fold), while 10 genes expressing different porins or their regulators were downregulated (∼2 - 3 fold). Except for porins, all these genes were further upregulated (∼2 – 6 fold) in swarm cells under antibiotic stress (Kan and Cip). The presence of AcrA also increased expression of these same genes (∼2 – 9 fold). 23 genes for energy metabolism (∼ 2 - 4 fold) and 16 genes related to ROS catabolism (∼ 2 - 6 fold) were also upregulated. **b**. Efflux assay using Nile red indicator dye. Glucose was added at 100s. Non-linear regression analysis of Planktonic, Planktonic+AcrA, Swarm, and Swarm+AcrA data sets (see d for color key) yielded rate constants of 1.001×10^−6^, 8.669×10^−007^, 6.630×10^−006^, and 1.165×10^−006^ respectively. A.U. = (A_636_/OD_600_). The time required for 50% Nile Red efflux or t_eff50,_ was significantly low in swarm cells (230s for swarm and ≥300s for planktonic cells; compare red vs green lines; p <0.0001 for the two values). Addition of AcrA yielded t_efflux50_ with a 1/10^th^ increase in swarm cells (207s, (p=0.0003 for Swarm+AcrA vs Swarm) and 1/20^th^ increase in planktonic cells (282s, compare brown vs blue lines; p=0.0038 for planktonic +AcrA vs Planktonic). **c**. Membrane permeability assay using Alamar Blue indicator dye. Non-linear regression analysis of Swarm, Swarm+AcrA, Swim, and Swim+ AcrA data sets (see d for color key) produced slopes of 0.06254, 0.06103, 0.02876, and 0.02498 respectively. A.U. = (A_590_/OD_600_**). d**. Estimated intracellular concentration of Fleroxacin. The slopes of the fitted lines of datasets for Swarm, Swarm+AcrA, Planktonic, and Planktonic+AcrA were 2.090, 1.220, 5.010, and 4.290 respectively. The keys for b-d are shown in d.

Taken together, the data indicate that swarm cells are intrinsically programmed to survive antibiotics by at least three different pathways: increased efflux, reduced permeability, and deployment of ROS catabolism (Fig. 4). Cell death (induced by antibiotic or other insults) further activates two of these pathways - efflux and ROS - by releasing AcrA, which binds to TolC on live cells (Fig. 2d) to instantly stimulate efflux (Fig. 3b), and to upregulate the expression of a multitude of genes for a sustained response (Fig. 3a). How AcrA triggers these events will be avenues of future research.

**Figure 4.**
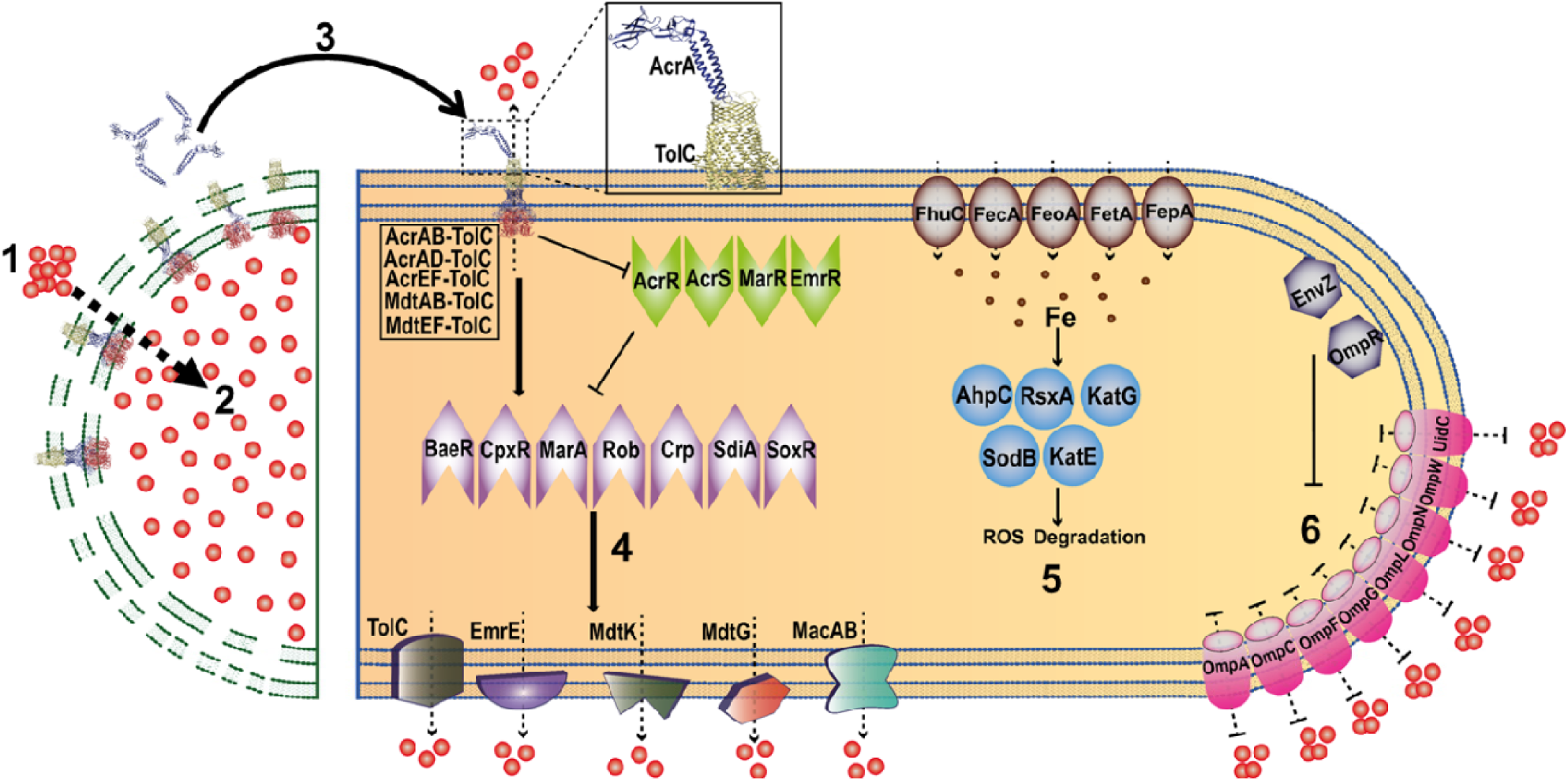
Model showing how AcrA necrosignaling promotes STRIVE. Both live and dead cells are represented in a single cartoon. **1**. Antibiotic uptake (red dots). **2**. Cell death, membrane damage. **3**. Released AcrA binds to TolC in the OM of live cells, activating efflux from TolC pumps. **4**. A secondary consequence of TolC activation is upregulation of five categories of efflux pumps via upregulation (arrowhead) and downregulation (flathead) of multiple transcription activators (purple) and repressors (green). **5**. Upregulation of genes that reduce ROS (blue and brown) would allow tolerance of the antibiotic stress. **6**. Downregulation (flathead) of porins (magenta) would restrict entry of antibiotics.

## Discussion

The surprising aspect of our findings is that a normal component of an RND drug efflux pump, AcrA, moonlights as a necrosignal directed at similar pumps. Another curious finding is the structural homology of AcrA to colicin E3 (Fig. S13), which interacts with BtuB (vitamin B12 receptor), a β-barrel OM protein similar to TolC^37,38^. We have used the colicin homology to suggest possible modes of AcrA binding to TolC (Fig. S14); the best fit has striking similarities with colicin E3-BtuB interaction^39^, suggesting a case of molecular mimicry. Comparison of AcrA from the bacterial species used in this study shows a high degree of sequence conservation (Fig. S15a, b), the *E. coli* and *Salmonella* proteins being most closely related (Fig. S15c), explaining their interchangeable necrosignaling response (Fig. 1e). Interestingly, TolC is absent in many bacteria including *Bacillus*. In contrast, AcrA is highly conseved across Bacterial domain (Fig. S16) suggesting it might have other binding partners, i.e., more necrosignaling modules may exist in nature. The existence of a specific signal-receptor survival module (AcrA:TolC) is expected to benefit its own species in a multi-species ecological community (Fig. 1e).

Death of a subpopulation for the overall survival of community and its resemblance to altruism, has been the subject of earlier studies^17,40,41^. While a variety of functions for cell lysis have been reported - recycling nutrients during bacterial starvation^42^, exchange of genetic material^43^, and activation of systems that deliver toxins to confer fitness in inter-species co-cultures^44^, the cell death response we report here have several features not shown in earlier studies. For example, the existence of a heterogeneous population with respect to antibiotic susceptibility (Figs. 1a and S1a), and the advantage it confers, is akin to a bet-hedging survival strategy^17^. High cell densities provide another selective advantage given that a large fraction of the swarm is killed for STRIVE to be observed (∼1/2 at Kan^25^; Fig. S1a), an observation that should be considered for clinical antibiotic regimen guidelines^45^ and in ecological studies involving microbial interactions^46,47^. A large motile population, that can continuously acquire new territory, and has a strategy to overcome stressful environments and survive mass extinction events promoted by the event itself, will also increase its chances of propagating beneficial mutations such as antibiotic resistance. We note that a response to cell death that benefits the living is not confined to bacteria, but is widespread in nature, as seen in insects (necromones)^48,49^, fishes^50^, birds^51^, and mammals^52^, possibly even in eukaryotic tumors^53^.

## Acknowledgements

This work was supported by National Institutes of Health Grant GM118085 and in part by the Robert Welch Foundation Grant F-1811. We thank Barrick lab (UT, Austin, Tx) for the ASKA library.

## Author contributions

SB and RMH set up the experimental design, SB performed the experiments, DMW modeled the survival data, SB and RMH wrote the manuscript.

## Methods

### Strains, plasmids and growth conditions

Bacterial strains used in this study: *Escherichia coli* (MG1655), *Salmonella enterica* serovar Typhimurium (14028), *Serratia marcescens* (274), *Bacillus subtilis* (3610), and *Pseudomonas aeruginosa* (PAO1). Oligonucleotide information will be provided upon request. The strains were propagated in LB (10 g/L tryptone, 5 g/L yeast extract, 5 g/L NaCl) broth or on 1.5% Bacto agar plates for single colony isolation. Antibiotics for marker selection were added as follows: Kan (Kanamycin) 25 μg/ml, Amp (Ampicillin) 100 μg/ml, Cam (Chloramphenicol) 30 μg/ml. Antibiotic concentrations are indicated henceforth (and in the text) with superscripts.

All gene deletions were achieved by transferring the Kan deletion marker from donor strains in the Keio collection^54^ to *E. coli* MG1655 using P1 transduction^55^ (P1*vir*). For overexpression analysis, genes were expressed from ASKA library^56^ pCA24N plasmid, induced with 0.1 mM IPTG. For AcrA purification, the periplasmic portion of AcrA without its lipoprotein signal peptide (as described^57^) was cloned in pTrc99a plasmid with an N- and C-terminal His and FLAG epitope tags, respectively. The *tolC* gene was cloned in pTrc99a, and site-directed mutagenesis was performed as described^58^.

### Motility assays

All assays used LB as the nutrient medium.

#### Swarm assays

Swarm plates were prepared with Bacto agar for all bacteria except for *E. coli* (which requires Eiken agar; Eiken Chem. Co. Japan) at the following agar concentrations: *E. coli* (0.5 %), *S. enterica* and *P. aeruginosa* (0.6%), *S. marcescens* (0.8%), and *B. subtilis* (0.7%). For *E. coli* and *S. enterica*, 0.5% glucose was included. Poured swarm plates were dried overnight at room temperature prior to use. Plates were incubated at 30°C for *E. coli* and 37°C for all others.

#### Border-crossing swarm assay

This was performed as described earlier^5^ and illustrated in Fig. S2. 4 μL of mid-log phase bacterial culture were inoculated in the left no-antibiotic chamber and allowed to dry by leaving the lid of the petri dish open for 15 minutes. The plates were then incubated at 30°C or 37°C for 16 h, which is the time it took for bacteria in control plates to colonize the entire no-antibiotic right chamber. All plates were photographed with a Canon Rebel XSI digital camera using the “bucket of light” as a light source^59^.

For border crossing assays in Fig. 1e, the right chamber contained Kan^50^ for *Ec, Se*, and *Bs*; Kan^75^ for *Sm*; Kan^125^ for *Pa*. Except for *Pseudomonas*, which was pre-killed by Kan^500^, all others were pre-killed Kan^250^.

### Preparation of dead cells and extracts

A 10-ml culture of *E. coli* cells (0.6 OD_600_) was treated for 30 min with any one of the following antibiotics: Kan^250^, Gen^50^, Amp^500^, and Cip^25^. Cells were harvested by centrifugation (12000 g, 3 min, 4°C). The pellet was once washed with sterile HEPES buffer (1.2 mM CaCl_2_, 1.2 mM MgCl_2_, 2.4 mM K_2_HPO_4_, 20 mM HEPES, 115 mM NaCl, pH of 7.4) and resuspended in the same buffer. Efficiency of killing was monitored by CFU counts on LB agar. 50 μl of the dead cell suspension was used for the border-crossing assay. To prepare extracts from the killed cells, the protocol above was scaled up to a 50 ml culture, the final cell pellet resuspended in 1 ml of HEPES buffer, and the cell suspension lysed by passing 3 times through a SLM AMINCO French press (1200 psig units). The lysed extract was precipitated by centrifugation (12000 g, 5 min, 4°C), and the supernatant used for further purification.

### Identification of the necrosignaling factor

The border-crossing assay was used to detect activity at all steps.

#### 1. Cell extract preparation

4 litre cultures (0.6 OD_600_) of *E. coli/S. enterica* cells were treated with Kan^250^ for 30 min. The cells were harvested by centrifugation (5000 g, 10 min, 4°C). The pellet was washed once with HEPES buffer (1.2 mM CaCl_2_, 1.2 mM MgCl_2_, 2.4 mM K_2_HPO_4_, 20 mM HEPES, 115 mM NaCl, one SIGMAFAST™ protease inhibitor tablet, pH of 7.4). The cell pellet (3.8 g) were resuspended in 15 ml of HEPES buffer and then lysed by six passages through a French Press (1200 psig units). The resultant extract was then centrifuged (10000 g for 10 min at 4°C) to collect the supernatant (50–70 mg/ml protein).

#### 2. Ammonium sulphate precipitation

The supernatant was treated with ammonium sulphate (AS) (30% for *E. coli* and 35% for *S. enterica*) overnight (O/N) in a cold room (4°C), while keeping it in a tumbling mode using a Barnstread thermolyne Labquake^TM^ rotisserie shaker. The precipitate was collected by centrifugation (12000 g for 30 min at 4°C), the pellet was resuspended in 5 ml of HEPES buffer and dialysed (O/N) using HEPES as the exchange buffer (∼5 mg/ml protein). 50 μl each of initial cell extract and the AS pellet were used to test for activity.

#### 3. Q sepharose anion exchange chromatography

3.5 ml of the AS fraction was loaded on a Q sepharose anion exchange chromatography column (10 cm × 0.8 cm^2^ = 8 ml), and after the flow-through was collected, the column was washed with HEPES buffer (1 column volume). The proteins were manually eluted (0.5 ml/min) using a 50-500 mM NaCl gradient in a total of fifteen-1ml fractions (∼2 column volumes). Fractions 5 and 6 showed activity for both strains. Their protein concentrations were ∼1.5 mg/ml.

#### 4. CM cation chromatography

The active fractions from step 3 were pooled and loaded onto a CM cation chromatographic column (7.5 cm × 0.8 cm^2^ = 6 ml). The flow-through was collected, followed up a 1-column volume wash with Phosphate-buffered saline (NaCl 137 mM, KCl 2.7 mM, Na_2_HPO_4_ 10 mM, KH2PO4 1.8 mM, pH 7.2). The proteins were manually eluted (0.5 ml/min) with a NaCl gradient of 50-250 mM using ∼2 column volumes (9 fractions×1.5 ml each=13.5 ml total). The first four fractions (50-125 mM NaCl) showed activity, and had protein concentrations in the range of 0.39 - 0.1 mg/ml.

### Mass spectrometric analysis

Active fractions from the CM cation step were used to perform ESI-MS in a Thermo Orbitrap Fusion hybrid mass spectrometer. The proteins were Trypsin digested on the column before Tandem mass spectra were extracted. All MS/MS samples were analyzed using Sequest (Thermo Fisher Scientific; version IseNode in Proteome Discoverer 1.4.1.14) and X! Tandem (CYCLONE 2010.12.01.1). Sequest and X! Tandem were searched with a fragment ion mass tolerance of 0.80 Da and a parent ion tolerance of 10.0 PPM. Carbamidomethyl of cysteine was specified in Sequest and X! Tandem as a fixed modification. Glu->pyro-Glu of the N-terminus, ammonia-loss of the N-terminus, Gln->pyro-Glu of the N-terminus and oxidation of methionine were specified in X! Tandem as variable modifications. Oxidation of methionine was specified in Sequest as a variable modification.

Scaffold (version Scaffold_4.8.4, Proteome Software Inc., Portland, OR) was used to validate MS/MS based peptide and protein identifications. Peptide identifications were accepted if they could be established at greater than 16.0% probability to achieve an FDR less than 1.0% by the Peptide Prophet algorithm^60^ with Scaffold delta-mass correction. Protein identifications were accepted if they could be established at greater than 95.0% probability to achieve an FDR less than 5.0% and contained at least 4 identified peptides. Protein probabilities were assigned by the Protein Prophet algorithm^61^. Proteins that contained similar peptides and could not be differentiated based on MS/MS analysis alone were grouped to satisfy the principles of parsimony.

### Purification of AcrA

BL21(DE3) harboring the His-AcrA-FLAG construct described above (refered to as simply AcrA henceforth) was cultured overnight in LB containing ampicillin^100^. An overnight culture was diluted 1:100 into 1 liter of fresh LB, and incubated at 37^0^C till 0.6 OD_600_ was reached. Protein Expression was induced with the addition of 1 mM IPTG for four hours. The cells were harvested by centrifugation, washed, resuspended in buffer [10 mM Tris-HCl (pH 8.0), 100 mM NaCl, one SIGMAFAST™ protease inhibitor tablet, 1 mg/ml lysozyme, 1 mM β-ME, 50 mM imidazole], and sonicated on ice. The supernatant was collected after centrifugation (12000g, 30 min, 4°C). Protein was purified using HisTrap HP (1 ml) column (GE) according to the manufacturer’s protocol with slight modifications. A 5-step gradient of imidazole (100 mM, 200mM, 300mM, 400mM, 500mM) was used to elute the bound protein manually (2 column volumes for each step) and 1ml fractions were collected. The first 6 fractions were pooled and dialyzed against two changes of buffer A (10 mM Tris-HCl (pH 8.0), 100 mM NaCl, 1 mM EDTA) overnight.

### RNA-Seq analysis

*E. coli* cells were grown in 10 ml broth (planktonic) culture (0.6 OD_600_) in absence of antibiotic and harvested with or without the treatment of antibiotic (see figure legends for specific experiments). *E. coli* swarm cells from the right chamber in the border-crossing assay (with or without antibiotics) were resuspended in 3 ml LB medium and harvested. The harvested cells were resuspended in 1 ml ice-chilled LB medium, kept on ice for 2 minutes, and pelleted by centrifugation (4°C, 10000g, 2 minutes). Every experimental condition tested had two biological replicates. Total RNA was isolated from these samples using Qiagen RNeasy® Protect Bacteria Mini Kit and following the enzymatic lysis method. The RNA was then used for library preparation using NEBNext Ultra II Library Prep Kit and sequenced on an Illumina NextSeq 500 platform (SR 75) yielding a total of 267.1 million reads for 10 samples. Sequence quality was determined by FastQC v0.13 and MultiQC^62^. The raw files were processed (Cutadapt^63^), aligned (Bowtie2^64^), mapped (Samtools^65^, Bedtools^66^) to reference genome (GenBank ID U00096.3), normalized and analyzed (DESeq2^67^), and visualized (R studio, Microsoft Excel, and Graphpad).

### Microscopy

The Kan marker from Δ*tolC* and Δ*acrA* strains were removed using a pCP20 based method^68^. To prepare the AcrA probe, DYKDDDDK Tag mouse monoclonal antibody (FG4R) (Thermo Fisher) was labelled with a quantum dot (Qdot 705) using a click chemistry kit (SiteClick™ Qdot™ 705 Antibody Labeling Kit, Thermo Fisher)^69^. 100 μl of either *E. coli* planktonic culture in LB medium (0.6 OD_600_, treated with Kan^25^ for 30 min) or swarm cells collected from border crossing assay plates (Kan^25^) in the same way as described in RNA seq section,) were incubated with membrane dye FM1-43 (0.1 mg/ml, FilmTracer™ FM™ 1-43, Thermo Fisher) for 15 minutes at 37°C in the dark. 100 ng of purified AcrA was added to these cell suspensions and incubated for 15 more minutes. The cells were then pelleted, washed, resuspended in 100 μl of 1x PBS (pH 7.4), and Qdot 705-labelled antibody was added (1:1000) for 15-minute incubation at room temperature. The excess antibody was removed by washing the cells by PBS buffer. Cells were then visualized using a light microscope (BX53F; Olympus, Tokyo, Japan) and cellSens software (v1.6) with minimal adjustments and alignments in Adobe Photoshop. The dye FM1-43 was visualized using a GFP filter (excitation wavelength, 460-480nm; emission 495-540nm). To reduce quenching and avoid bleed-through because of overlapping excitation or emission wavelengths, the Qdot705 was visualized using a bandpass filter AT-Qdot 705 filter (Chroma, 39018) that allows excitation wavelength of 400-450 nm (425 CWL) and emission wavelength of 685-725 nm (705 CWL). After incubation with AcrA for the microscopy experiments, a subset of the swarm cell sample (200 μl) was used to determine the localization of extracellularly added AcrA as described^70^ with modifications. The cells were pelleted with gentle centrifugation (1500 g, 10 min, 4°C) and resuspended in protease buffer (50 mM Tris, 7.5 mM CaCl_2_, pH 8.8). The suspension was divided into two equal halves. One half was lysed with sonication (three cycles of 30 s on and 30 s off) and supernatant was collected. The un-lysed cells and the supernatant of the lysed sample were incubated with trypsin (50 μg/ml, Thermo Fisher) for 30 min at RT. The protease was then inactivated with 0.1 M PMSF. These samples were used for western blot analysis with DnaK as cytoplasmic control using mouse monoclonal anti-DnaK antibody (abcam).

### Efflux pump assays

Planktonic and swarm cells were harvested as for RNA seq analysis. Nile Red and Alamar Blue assays were performed as described^71^ with modifications. Each sample (1 mL) was centrifuged for 10 min at 5000g; the pellet was suspended in PPB (20 mM potassium phosphate, 1 mM MgCl_2_, pH 7.0) to obtain OD_600_ of 1.0.

#### Nile Red assay

Samples were incubated with 10 mM CCCP (prepared in 50% DMSO) for 30 min at RT followed by addition of 10 mM Nile Red (VWR) at 37°C and shaken at 200 rpm for 30 min. These cells were then kept in room temperature for 15 min without shaking, harvested via centrifugation, and supernatant was carefully and completely discarded. The resultant pellet was resuspended in PPB to obtain OD_600_ of ∼1.0; a 10x dilution of that suspension was transferred to a quartz cuvette to measure fluorescence (excitation at 552 and emission at 636 nm) using a QuantaMaster spectrofluorometer every 10 s for 100s with intermittent manual stirring. The efflux of Nile Red was triggered by addition of 100 μl of 1 M glucose followed by measuring fluorescence every 10 s for 200s. In a subset of samples, 100 ng of purified AcrA was added along with the glucose. The obtained absorbance (A_636_) was normalized with OD_600_.

#### Alamar Blue assay

20 μl of Alamar Blue liquid reagent (Invitrogen) was added to 180 μl of cell suspension. The fluorescence of the samples was measured (excitation at 565nm, emission at 590nm) every 5 min for 30 min followed by every 10 min for 30 more min. In a subset of samples, 20 ng of purified AcrA was added before the addition of Alamar Blue.

#### Fleroxacin assay^72^

Planktonic and swarm cells were harvested as before. A cell suspension of ∼0.6 OD_600_ were prepared in 2 ml LB and each sample was divided into two equal halves. Appropriate amount of Fleroxacin (VWR) were added to one-half of the cells in media and the cells were incubated for 30 min in shaking condition at 37°C. The other half was used as the untreated control (990 μl) and to calculate CFU (10 μl). The cells were pelleted, washed once, resuspended in 1 ml of 50 mM sodium phosphate buffer (pH 7.2), and sonicated to lyse (3 rounds of 30s on and 30s off). The supernatant was used to determine fluorescence at 442 nm with excitation at 282 nm. The actual Fleroxacin fluorescence was calculated by subtracting the fluorescence of untreated sample from fluorescence of treated samples. The concentration of Fleroxacin was determined by comparing with a standard curve of known Fleroxacin concentration.

#### Disc diffusion assay

100 μl of collected cells (as described in supplementary figure S11) were kept at 95°C for 15 minutes, chilled in 4°C for 5 minutes, and centrifuged at 10000g for 5 min. The supernatant was used to deposit on discs. Kan is a thermostable antibiotic^34^.

### Killing curves

An O/N culture of *E. coli* was sub-cultured (1 ml) to obtain 0.6 OD_600_. The swarm cells were collected from a regular swarm plate using 1 ml of LB and finally diluted to 0.6 OD_600_. These cell suspensions were then incubated with various concentrations of Kan for 2 hours at 37°C with shaking at 200 rpm. A fraction of cells (100 μl) were collected every 30 min and dilution plating was used for measuring CFU counts. To confirm CFU counts, another fraction of cells (100 μl) from some samples were stained with LIVE/DEAD Bac Light Bacterial Viability Kit (Thermo Fisher) for 15 min by adding 5 μl of SYTO9 and PI dye each. The cells were visualized using a fluorescent microscope (BX53F; Olympus, Tokyo, Japan) with GFP and RFP filters for SYTO9 and PI dyes respectively.

### External AcrA docking on TolC

AcrA and TolC structures were obtained from PDB^73^ (ID: 5NG5)(rcsb.org). One AcrA molecule was docked on to TolC tripartite complex using the HADDOCK2.2 web server^74^ (https://haddock.science.uu.nl/services/HADDOCK2.2/) with default parameters The docking was guided excluding the the region of TolC that normally interacts with AcrA in the periplasm (Gln142-Thr152 and Ala360-Val372), but the whole AcrA structure was used.The resulting top four models had HADDOCK scores ranging from -172 to -139, z-scores from -2.5 to -0.4, and RMSD of 0.7 to 2.3Å.

### AcrA sequence comparison and distribution

AcrA sequences from five organisms studied here were used for multiple sequence alignment (MSA) using Clustal Omega^75^ algorithm and visualized by Jalview^76^. The sequences were obtained from NCBI protein database (https://www.ncbi.nlm.nih.gov/protein). A cladogram was generated by neighbor-joining method without distance corrections using Clustal Omega showing relatedness of these AcrA sequences.

The *E. coli* AcrA and TolC sequences were separately used as inputs in the homology-based protein sequence alignment algorithm phmmer in HMMER^77^ web server (https://www.ebi.ac.uk/Tools/hmmer/) with a significant e-value and hit cutoffs of 10^−20^ and 0.03, respectively. The hits were then visualized in HMMER taxonomy and counts for each taxon in bacterial domain were used for generating a heat map (Fig. S16).

## References

1 Kearns, D. B. A field guide to bacterial swarming motility. Nature reviews. Microbiology 8, 634–644, doi:10.1038/nrmicro2405 (2010).

2 Kim, W., Killam, T., Sood, V. & Surette, M. G. Swarm-cell differentiation in Salmonella enterica serovar typhimurium results in elevated resistance to multiple antibiotics. Journal of bacteriology 185, 3111–3117 (2003).

3 Overhage, J., Bains, M., Brazas, M. D. & Hancock, R. E. Swarming of Pseudomonas aeruginosa is a complex adaptation leading to increased production of virulence factors and antibiotic resistance. Journal of bacteriology 190, 2671–2679, doi:10.1128/JB.01659-07 (2008).

4 Lai, S., Tremblay, J. & Deziel, E. Swarming motility: a multicellular behaviour conferring antimicrobial resistance. Environmental microbiology 11, 126–136, doi:10.1111/j.1462-2920.2008.01747.x (2009).

5 Butler, M. T., Wang, Q. & Harshey, R. M. Cell density and mobility protect swarming bacteria against antibiotics. Proceedings of the National Academy of Sciences of the United States of America 107, 3776–3781, doi:10.1073/pnas.0910934107 (2010).

6 Piddock, L. J. Multidrug-resistance efflux pumps - not just for resistance. Nature reviews. Microbiology 4, 629–636, doi:10.1038/nrmicro1464 (2006).

7 Du, D. et al. Multidrug efflux pumps: structure, function and regulation. Nature reviews. Microbiology 16, 523–539, doi:10.1038/s41579-018-0048-6 (2018).

8 Sandoval-Motta, S. & Aldana, M. Adaptive resistance to antibiotics in bacteria: a systems biology perspective. Wiley interdisciplinary reviews. Systems biology and medicine 8, 253–267, doi:10.1002/wsbm.1335 (2016).

9 Jarrell, K. F. & McBride, M. J. The surprisingly diverse ways that prokaryotes move. Nature reviews. Microbiology 6, 466–476, doi:10.1038/nrmicro1900 (2008).

10 Mazzantini, D. et al. FlhF Is Required for Swarming Motility and Full Pathogenicity of Bacillus cereus. Frontiers in microbiology 7, 1644, doi:10.3389/fmicb.2016.01644 (2016).

11 Mobley, H. L. & Belas, R. Swarming and pathogenicity of Proteus mirabilis in the urinary tract. Trends in microbiology 3, 280–284, doi:10.1016/s0966-842x(00)88945-3 (1995).

12 Kim, W. & Surette, M. G. Swarming populations of Salmonella represent a unique physiological state coupled to multiple mechanisms of antibiotic resistance. Biological procedures online 5, 189–196, doi:10.1251/bpo61 (2003).

13 Stewart, P. S. Mechanisms of antibiotic resistance in bacterial biofilms. International journal of medical microbiology : IJMM 292, 107–113, doi:10.1078/1438-4221-00196 (2002).

14 Balaban, N. Q. et al. Definitions and guidelines for research on antibiotic persistence. Nature reviews. Microbiology 17, 441–448, doi:10.1038/s41579-019-0196-3 (2019).

15 Brauner, A., Fridman, O., Gefen, O. & Balaban, N. Q. Distinguishing between resistance, tolerance and persistence to antibiotic treatment. Nature reviews. Microbiology 14, 320–330, doi:10.1038/nrmicro.2016.34 (2016).

16 Partridge, J. D., Ariel, G., Schvartz, O., Harshey, R. M. & Be’er, A. The 3D architecture of a bacterial swarm has implications for antibiotic tolerance. Scientific reports 8, 15823, doi:10.1038/s41598-018-34192-2 (2018).

17 Allocati, N., Masulli, M., Di Ilio, C. & De Laurenzi, V. Die for the community: an overview of programmed cell death in bacteria. Cell death & disease 6, e1609, doi:10.1038/cddis.2014.570 (2015).

18 Rice, K. C. & Bayles, K. W. Molecular control of bacterial death and lysis. Microbiology and molecular biology reviews : MMBR 72, 85-109, table of contents, doi:10.1128/MMBR.00030-07 (2008).

19 Ackermann, M. et al. Self-destructive cooperation mediated by phenotypic noise. Nature 454, 987–990, doi:10.1038/nature07067 (2008).

20 Anes, J., McCusker, M. P., Fanning, S. & Martins, M. The ins and outs of RND efflux pumps in Escherichia coli. Frontiers in microbiology 6, 587, doi:10.3389/fmicb.2015.00587 (2015).

21 Hulme, E. C. & Trevethick, M. A. Ligand binding assays at equilibrium: validation and interpretation. British journal of pharmacology 161, 1219–1237, doi:10.1111/j.1476-5381.2009.00604.x (2010).

22 Du, D. et al. Structure of the AcrAB-TolC multidrug efflux pump. Nature 509, 512–515, doi:10.1038/nature13205 (2014).

23 Zakharov, S. D., Wang, X. S. & Cramer, W. A. The Colicin E1 TolC-Binding Conformer: Pillar or Pore Function of TolC in Colicin Import? Biochemistry 55, 5084–5094, doi:10.1021/acs.biochem.6b00621 (2016).

24 German, G. J. & Misra, R. The TolC protein of Escherichia coli serves as a cell-surface receptor for the newly characterized TLS bacteriophage. J Mol Biol 308, 579–585, doi:10.1006/jmbi.2001.4578 (2001).

25 Stewart, B. J. & McCarter, L. L. Lateral flagellar gene system of Vibrio parahaemolyticus. Journal of bacteriology 185, 4508–4518, doi:10.1128/jb.185.15.4508-4518.2003 (2003).

26 Wang, Q., Frye, J. G., McClelland, M. & Harshey, R. M. Gene expression patterns during swarming in Salmonella typhimurium: genes specific to surface growth and putative new motility and pathogenicity genes. Molecular microbiology 52, 169–187, doi:10.1111/j.1365-2958.2003.03977.x (2004).

27 Kim, W. & Surette, M. G. Metabolic differentiation in actively swarming Salmonella. Molecular microbiology 54, 702–714, doi:10.1111/j.1365-2958.2004.04295.x (2004).

28 Dwyer, D. J. et al. Antibiotics induce redox-related physiological alterations as part of their lethality. Proceedings of the National Academy of Sciences of the United States of America 111, E2100–2109, doi:10.1073/pnas.1401876111 (2014).

29 Kohanski, M. A., Dwyer, D. J., Hayete, B., Lawrence, C. A. & Collins, J. J. A common mechanism of cellular death induced by bactericidal antibiotics. Cell 130, 797–810, doi:10.1016/j.cell.2007.06.049 (2007).

30 Masuda, I. et al. tRNA Methylation Is a Global Determinant of Bacterial Multi-drug Resistance. Cell systems 8, 475, doi:10.1016/j.cels.2019.05.002 (2019).

31 Bohnert, J. A., Karamian, B. & Nikaido, H. Optimized Nile Red efflux assay of AcrAB-TolC multidrug efflux system shows competition between substrates. Antimicrobial agents and chemotherapy 54, 3770–3775, doi:10.1128/AAC.00620-10 (2010).

32 Rampersad, S. N. Multiple applications of Alamar Blue as an indicator of metabolic function and cellular health in cell viability bioassays. Sensors 12, 12347–12360, doi:10.3390/s120912347 (2012).

33 Cinquin, B. et al. Microspectrometric insights on the uptake of antibiotics at the single bacterial cell level. Scientific reports 5, 17968, doi:10.1038/srep17968 (2015).

34 Traub, W. H. & Leonhard, B. Heat stability of the antimicrobial activity of sixty-two antibacterial agents. The Journal of antimicrobial chemotherapy 35, 149–154, doi:10.1093/jac/35.1.149 (1995).

35 Baker, R. D., Cook, C. O. & Goodwin, D. C. Properties of catalase-peroxidase lacking its C-terminal domain. Biochemical and biophysical research communications 320, 833–839, doi:10.1016/j.bbrc.2004.06.026 (2004).

36 Inoue, T. et al. Genome-wide screening of genes required for swarming motility in Escherichia coli K-12. Journal of bacteriology 189, 950–957, doi:10.1128/JB.01294-06 (2007).

37 Cramer, W. A., Sharma, O. & Zakharov, S. D. On mechanisms of colicin import: the outer membrane quandary. The Biochemical journal 475, 3903–3915, doi:10.1042/BCJ20180477 (2018).

38 Jakes, K. S. The Colicin E1 TolC Box: Identification of a Domain Required for Colicin E1 Cytotoxicity and TolC Binding. Journal of bacteriology 199, doi:10.1128/JB.00412-16 (2017).

39 Kurisu, G. et al. The structure of BtuB with bound colicin E3 R-domain implies a translocon. Nature structural biology 10, 948–954, doi:10.1038/nsb997 (2003).

40 Bayles, K. W. Bacterial programmed cell death: making sense of a paradox. Nature reviews. Microbiology 12, 63–69, doi:10.1038/nrmicro3136 (2014).

41 Nedelcu, A. M., Driscoll, W. W., Durand, P. M., Herron, M. D. & Rashidi, A. On the paradigm of altruistic suicide in the unicellular world. Evolution; international journal of organic evolution 65, 3–20, doi:10.1111/j.1558-5646.2010.01103.x (2011).

42 Takano, S., Pawlowska, B. J., Gudelj, I., Yomo, T. & Tsuru, S. Density-Dependent Recycling Promotes the Long-Term Survival of Bacterial Populations during Periods of Starvation. mBio 8, doi:10.1128/mBio.02336-16 (2017).

43 Takeuchi, N., Kaneko, K. & Koonin, E. V. Horizontal gene transfer can rescue prokaryotes from Muller’s ratchet: benefit of DNA from dead cells and population subdivision. G3 4, 325–339, doi:10.1534/g3.113.009845 (2014).

44 LeRoux, M. et al. Kin cell lysis is a danger signal that activates antibacterial pathways of Pseudomonas aeruginosa. eLife 4, doi:10.7554/eLife.05701 (2015).

45 Aidara-Kane, A. et al. World Health Organization (WHO) guidelines on use of medically important antimicrobials in food-producing animals. Antimicrobial resistance and infection control 7, 7, doi:10.1186/s13756-017-0294-9 (2018).

46 Ghoul, M. & Mitri, S. The Ecology and Evolution of Microbial Competition. Trends in microbiology 24, 833–845, doi:10.1016/j.tim.2016.06.011 (2016).

47 Tecon, R., Ebrahimi, A., Kleyer, H., Erev Levi, S. & Or, D. Cell-to-cell bacterial interactions promoted by drier conditions on soil surfaces. Proceedings of the National Academy of Sciences of the United States of America 115, 9791–9796, doi:10.1073/pnas.1808274115 (2018).

48 Sun, Q., Haynes, K. F. & Zhou, X. Managing the risks and rewards of death in eusocial insects. Philosophical transactions of the Royal Society of London. Series B, Biological sciences 373, doi:10.1098/rstb.2017.0258 (2018).

49 McAfee, A. et al. A death pheromone, oleic acid, triggers hygienic behavior in honey bees (Apis mellifera L.). Scientific reports 8, 5719, doi:10.1038/s41598-018-24054-2 (2018).

50 Hussain, A. et al. High-affinity olfactory receptor for the death-associated odor cadaverine. Proceedings of the National Academy of Sciences of the United States of America 110, 19579–19584, doi:10.1073/pnas.1318596110 (2013).

51 Swift, K. & Marzluff, J. M. Occurrence and variability of tactile interactions between wild American crows and dead conspecifics. Philosophical transactions of the Royal Society of London. Series B, Biological sciences 373, doi:10.1098/rstb.2017.0259 (2018).

52 Prounis, G. S. & Shields, W. M. Necrophobic behavior in small mammals. Behavioural processes 94, 41–44, doi:10.1016/j.beproc.2012.12.001 (2013).

53 Garg, A. D., Martin, S., Golab, J. & Agostinis, P. Danger signalling during cancer cell death: origins, plasticity and regulation. Cell death and differentiation 21, 26–38, doi:10.1038/cdd.2013.48 (2014).

54 Baba, T. et al. Construction of Escherichia coli K-12 in-frame, single-gene knockout mutants: the Keio collection. Molecular systems biology 2, 2006 0008, doi:10.1038/msb4100050 (2006).

55 Sternberg, N. L. & Maurer, R. Bacteriophage-mediated generalized transduction in Escherichia coli and Salmonella typhimurium. Methods in enzymology 204, 18–43, doi:10.1016/0076-6879(91)04004-8 (1991).

56 Kitagawa, M. et al. Complete set of ORF clones of Escherichia coli ASKA library (a complete set of E. coli K-12 ORF archive): unique resources for biological research. DNA research : an international journal for rapid publication of reports on genes and genomes 12, 291–299, doi:10.1093/dnares/dsi012 (2005).

57 Zgurskaya, H. I. & Nikaido, H. AcrA is a highly asymmetric protein capable of spanning the periplasm. Journal of molecular biology 285, 409–420, doi:10.1006/jmbi.1998.2313 (1999).

58 Liu, H. & Naismith, J. H. An efficient one-step site-directed deletion, insertion, single and multiple-site plasmid mutagenesis protocol. BMC biotechnology 8, 91, doi:10.1186/1472-6750-8-91 (2008).

59 Parkinson, J. S. A “bucket of light” for viewing bacterial colonies in soft agar. Methods in enzymology 423, 432–435, doi:10.1016/S0076-6879(07)23020-4 (2007).

60 Keller, A., Nesvizhskii, A. I., Kolker, E. & Aebersold, R. Empirical statistical model to estimate the accuracy of peptide identifications made by MS/MS and database search. Analytical chemistry 74, 5383–5392 (2002).

61 Nesvizhskii, A. I., Keller, A., Kolker, E. & Aebersold, R. A statistical model for identifying proteins by tandem mass spectrometry. Analytical chemistry 75, 4646–4658 (2003).

62 Ewels, P., Magnusson, M., Lundin, S. & Kaller, M. MultiQC: summarize analysis results for multiple tools and samples in a single report. Bioinformatics 32, 3047–3048, doi:10.1093/bioinformatics/btw354 (2016).

63 Martin, M. Cutadapt removes adapter sequences from high-throughput sequencing reads. EMBnet.journal 17, 10–12 (2011).

64 Langmead, B. & Salzberg, S. L. Fast gapped-read alignment with Bowtie 2. Nature methods 9, 357–359, doi:10.1038/nmeth.1923 (2012).

65 Li, H. et al. The Sequence Alignment/Map format and SAMtools. Bioinformatics 25, 2078–2079, doi:10.1093/bioinformatics/btp352 (2009).

66 Quinlan, A. R. & Hall, I. M. BEDTools: a flexible suite of utilities for comparing genomic features. Bioinformatics 26, 841–842, doi:10.1093/bioinformatics/btq033 (2010).

67 Love, M. I., Huber, W. & Anders, S. Moderated estimation of fold change and dispersion for RNA-seq data with DESeq2. Genome biology 15, 550, doi:10.1186/s13059-014-0550-8 (2014).

68 Datsenko, K. A. & Wanner, B. L. One-step inactivation of chromosomal genes in Escherichia coli K-12 using PCR products. Proceedings of the National Academy of Sciences of the United States of America 97, 6640–6645, doi:10.1073/pnas.120163297 (2000).

69 Zeglis, B. M. et al. Enzyme-mediated methodology for the site-specific radiolabeling of antibodies based on catalyst-free click chemistry. Bioconjugate chemistry 24, 1057–1067, doi:10.1021/bc400122c (2013).

70 Besingi, R. N. & Clark, P. L. Extracellular protease digestion to evaluate membrane protein cell surface localization. Nature protocols 10, 2074–2080, doi:10.1038/nprot.2015.131 (2015).

71 Masuda, I. et al. tRNA Methylation Is a Global Determinant of Bacterial Multi-drug Resistance. Cell systems 8, 302–314 e308, doi:10.1016/j.cels.2019.03.008 (2019).

72 Chapman, J. S. & Georgopapadakou, N. H. Fluorometric assay for fleroxacin uptake by bacterial cells. Antimicrobial agents and chemotherapy 33, 27–29, doi:10.1128/aac.33.1.27 (1989).

73 Burley, S. K. et al. RCSB Protein Data Bank: biological macromolecular structures enabling research and education in fundamental biology, biomedicine, biotechnology and energy. Nucleic acids research 47, D464–D474, doi:10.1093/nar/gky1004 (2019).

74 van Zundert, G. C. P. et al. The HADDOCK2.2 Web Server: User-Friendly Integrative Modeling of Biomolecular Complexes. Journal of molecular biology 428, 720–725, doi:10.1016/j.jmb.2015.09.014 (2016).

75 Sievers, F. et al. Fast, scalable generation of high-quality protein multiple sequence alignments using Clustal Omega. Molecular systems biology 7, 539, doi:10.1038/msb.2011.75 (2011).

76 Waterhouse, A. M., Procter, J. B., Martin, D. M., Clamp, M. & Barton, G. J. Jalview Version 2--a multiple sequence alignment editor and analysis workbench. Bioinformatics 25, 1189–1191, doi:10.1093/bioinformatics/btp033 (2009).

77 Finn, R. D. et al. HMMER web server: 2015 update. Nucleic acids research 43, W30–38, doi:10.1093/nar/gkv397 (2015).

