## Supplementary Figures and Text for "Necrosignaling: Cell death triggers antibiotic survival pathways in bacterial swarms"

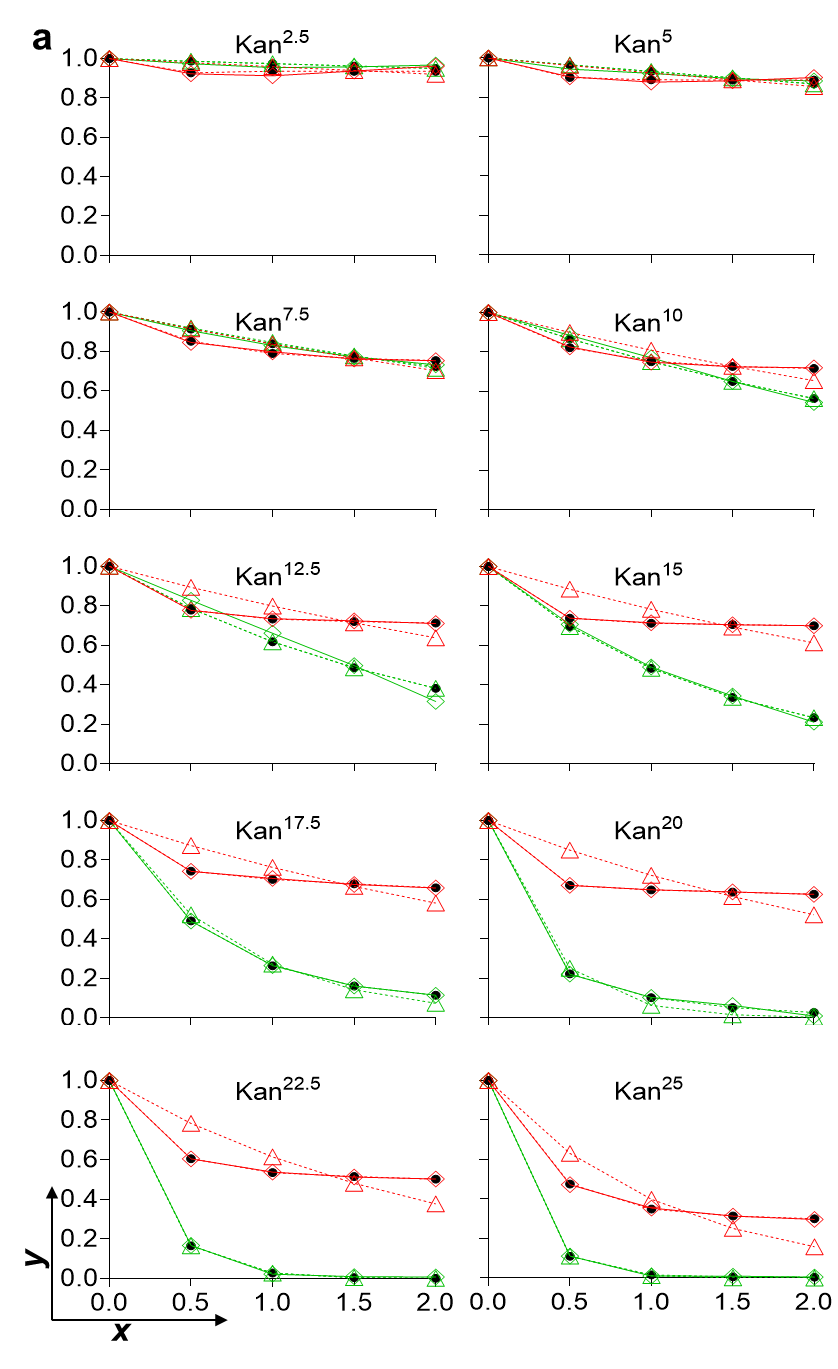

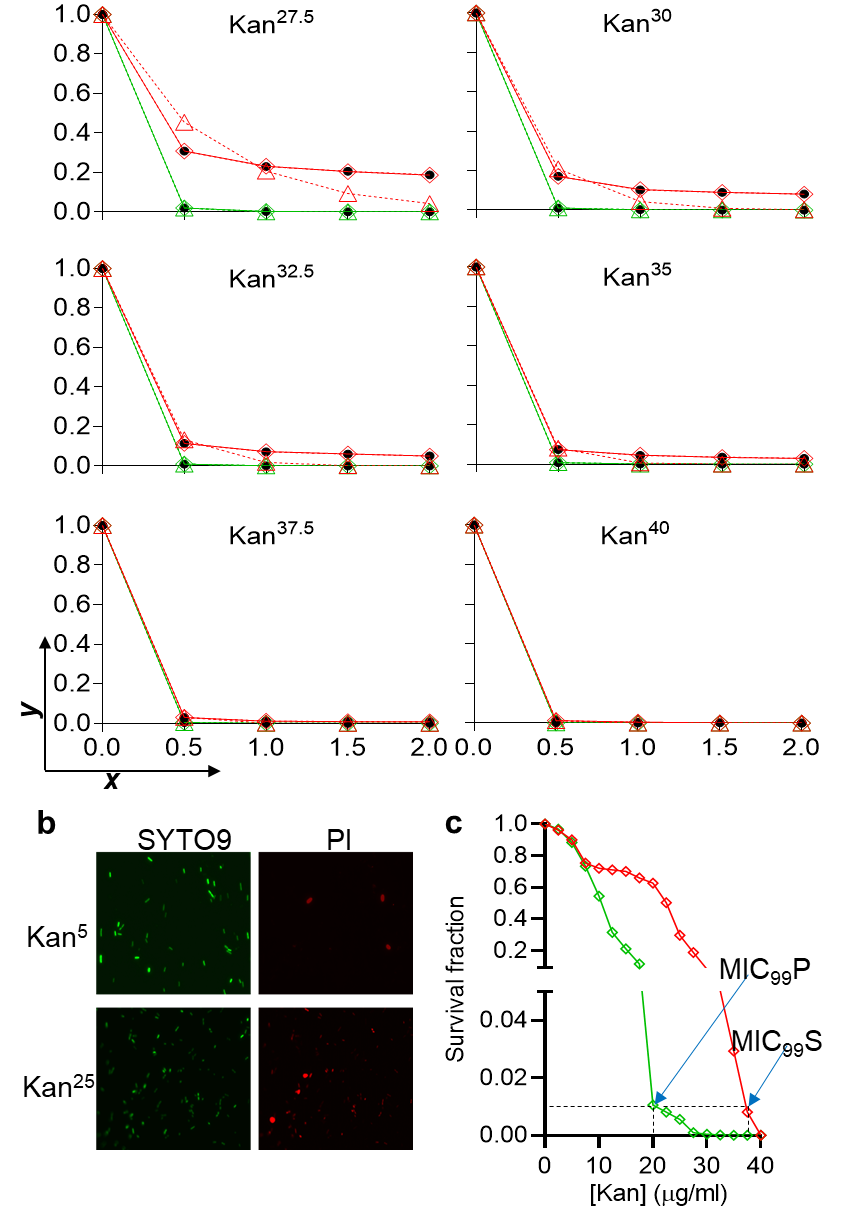

**Figure S1 |** **Experimental and simulated survival curves of swarm vs planktonic cells at varying kanamycin (Kan) concentrations.** **a.** Survival curves at Kan concentrations ranging from 2.5 to 40 μg/ml, showing our key observation that the best fit of the experimental data is to simulations that assume a heterogeneous swarm population and homogeneous planktonic population. Time in hours is plotted on the ***x*** axis, and fraction survived (as determined by CFU counts) on the ***y*** axis. The superscript following Kan refers to μg/ml. Solid red and green lines report experimental data for swarm and planktonic cells, respectively. Two sets of red and green dotted lines are simulated fits to the experimental data assuming either a homogenous (triangles) or heterogenous (circles) population of these two cells types, as described in Supplemental Text 1. **b.** Representative Live-Dead staining images of swarm cells treated with Kan^5^ and Kan^25^ for 0.5 h, confirming the CFU count data in A. All the cells are stained with the green dye SYTO9, but only cells with membrane damage (dead cells) get stained with red dye propidium iodide (PI). **c.** Determination of MIC (minimum inhibitory concentration) for Kan from survival curves at 2 h shown in A. MIC_99_, the antibiotic concentration required to kill 99% of the population is Kan^19^ for planktonic cells (green, MIC_99_P), and Kan^37.5^ for swarm cells (red, MIC_99_S).

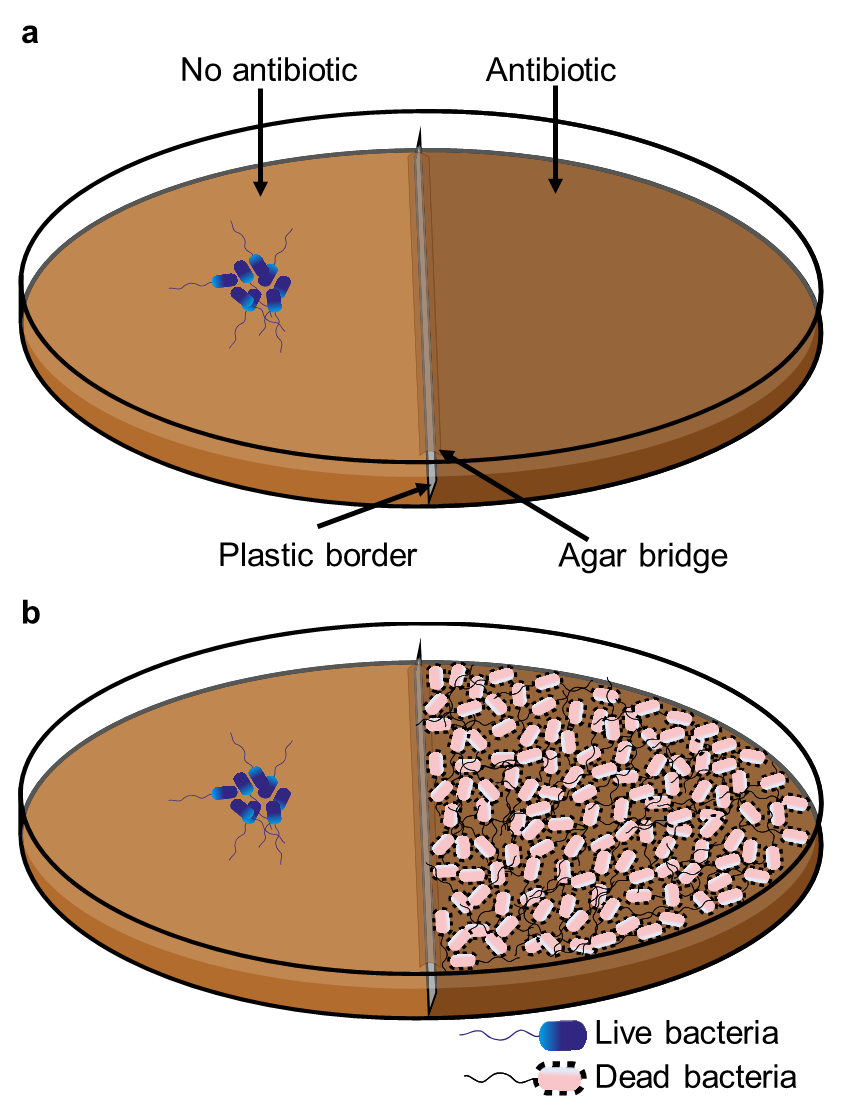

**Figure S2 |** **Schematic of the border-crossing assay.** **a**. Petri plates with a plastic divider create two chambers. The left chamber is poured with media without antibiotic, and the right with antibiotic. After the media is set, the two chambers are connected by a thin layer of agar on the top of the bridge as described^1^. Bacteria are inoculated in the left chamber as indicated, and allowed to swarm to the right chamber. **b**. As in A, but with dead bacteria layered on the surface of media on right as indicated.

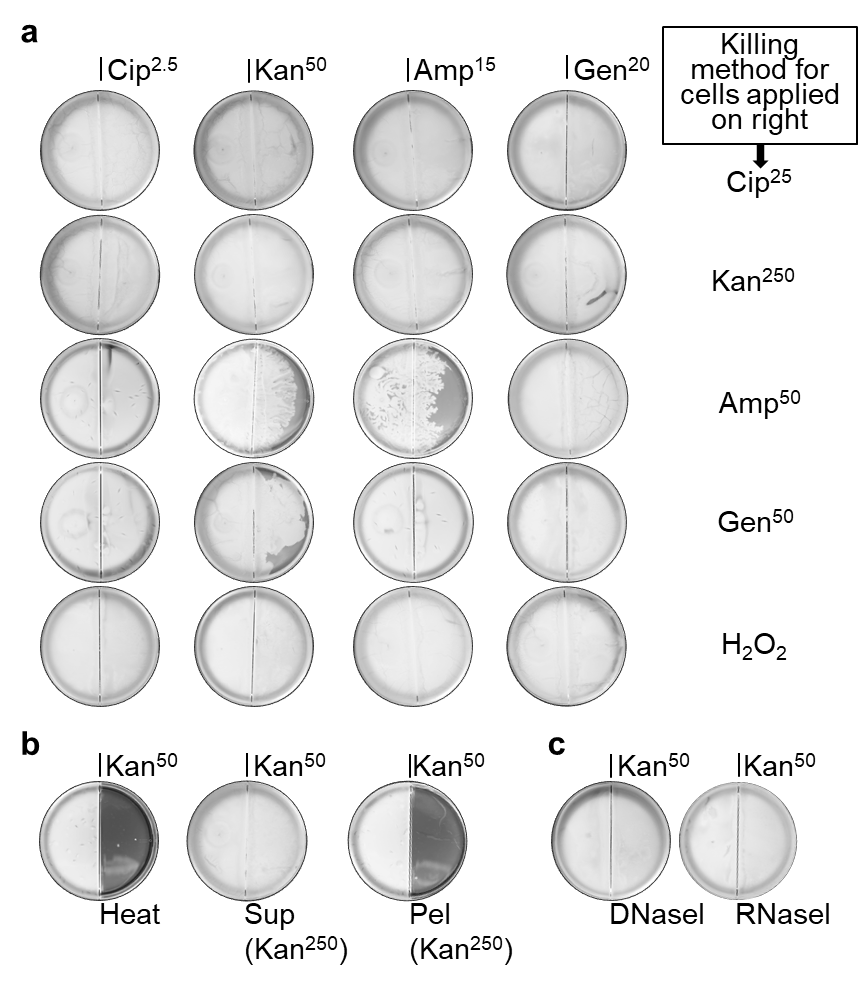

**Figure S3 | Dead cell-promoted STRIVE is independent of the killing method: necrosignaling factor is heat-sensitive and found in supernatants of cell extracts prepared from killed *E. coli* cells.** **a.** *E. coli* swarms are sensitive to Cip^2.5^, Kan^25^, Amp^15^ and Gen^20^. When cells killed by indicated methods are applied on the right, wild type *E. coli* cells inoculated on the left can migrate over the indicated antibiotic concentrations. **b.** Ability of heat killed cells, or of the supernatant (Sup) and pellet (Pel) fractions prepared from cells killed with Kan^250^, to support swarming on Kan^50^. **c**. Treatment of the Kan^250^ Sup with DNase and RNase does not destroy its necrosignaling activity.

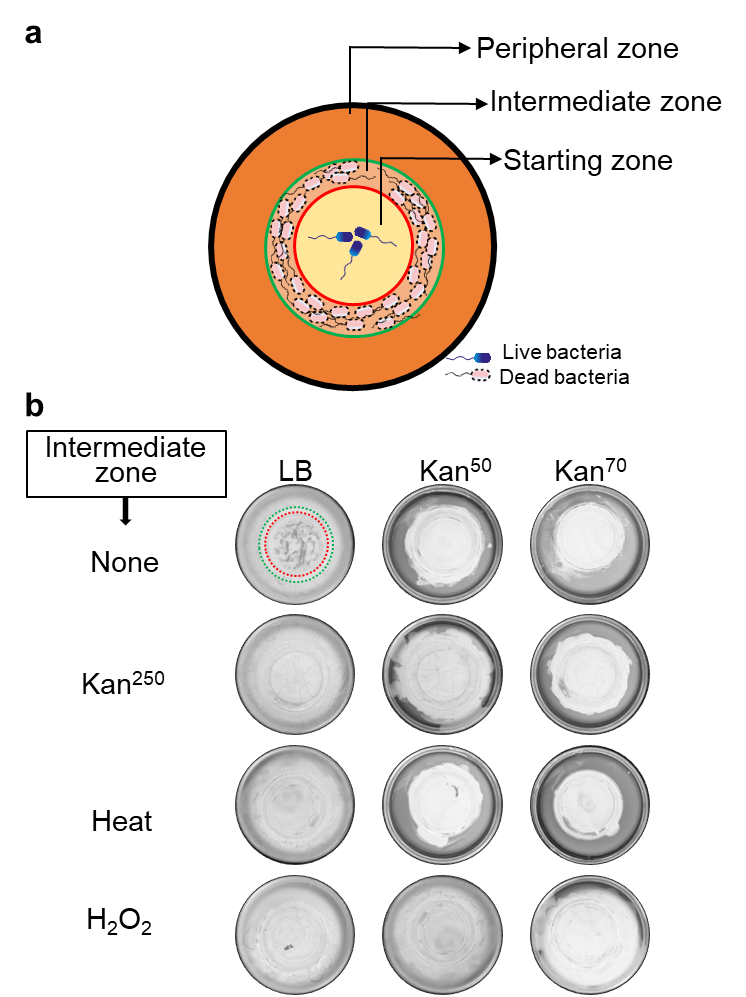

**Figure S4 | Tri-plate assay showing a sustained STRIVE response.** **a.** Cartoon demonstrating placement of plates of different diameters inside one another to create three swarming zones as labeled, with narrow agar bridges connecting the chambers as described in Fig. S2a. **b.** Wild-type *E. coli* was inoculated in the central zone. The composition of cells deposited on the intermediate zone is indicated on the left. The data show that cells swarming over the dead cell zone are potentiated for STRIVE even after exiting this zone, with 50 mM H_2_O_2_ eliciting the strongest response (promoting swarming on Kan^70^ in the peripheral zone). Dotted red and green circles on the top left tri-plate shows outlines of smallest and medium sized plates respectively.

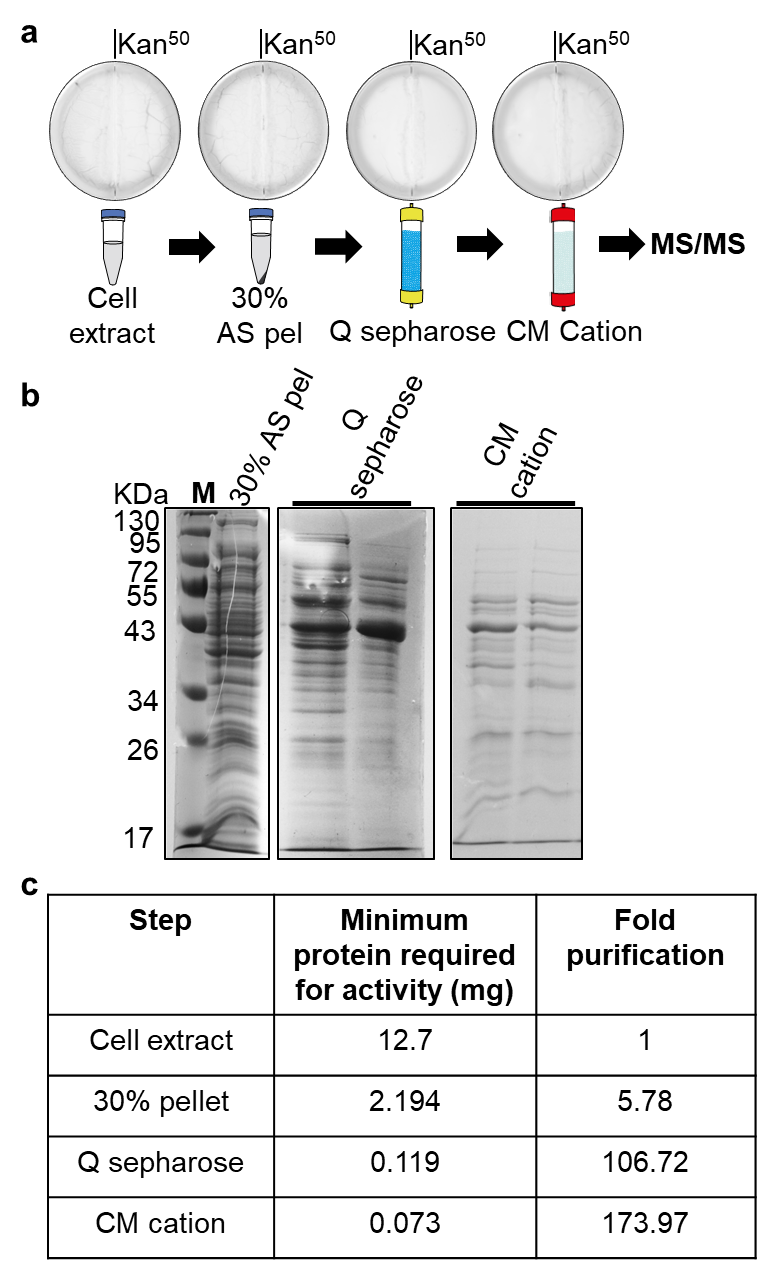

**Figure S5 |** **Purification of the necrosignal from *E. coli* and *Salmonella*.** **a**. Necrosignaling activity in cell extract supernatants was precipitated with ammonium sulfate (AS pel), resuspended and fractionated over Q Sepharose and CM Cation exchange columns, and analysed by MS/MS. See Methods for purification details. Plates shown are for *E. coli.* **b.** 12% SDS-PAGE gels showing *E. coli* active fractions from 30% AS pel, Q Sepharose (fractions 5 and 6) and CM cation (fractions 1 and 2); M, protein MW marker lane. **c**. Table showing fold-purification achieved in *E. coli*.

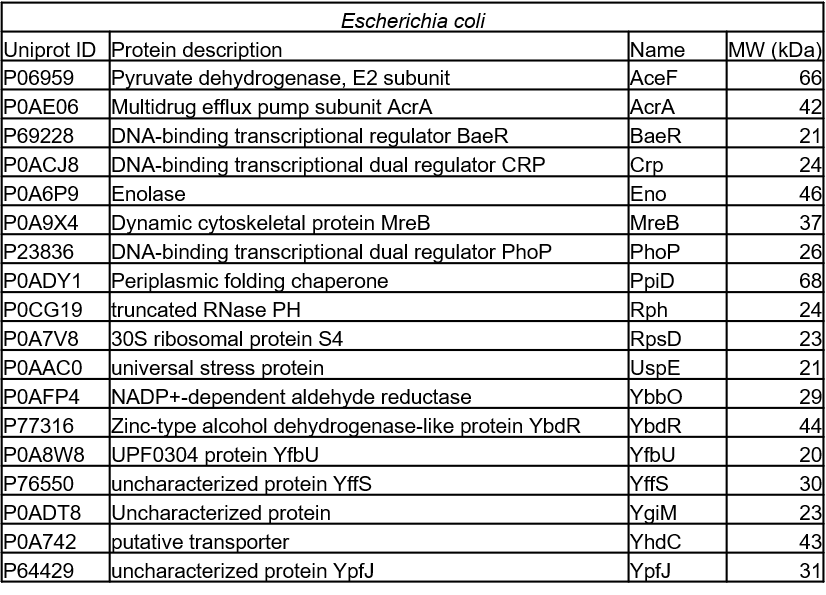

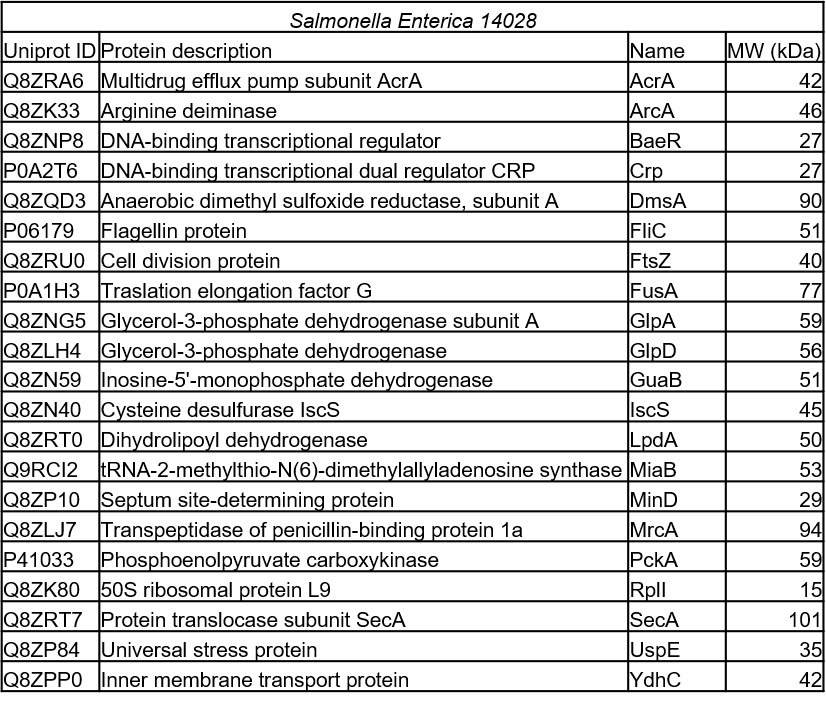

**Figure S6 | MS/MS analysis of the SR-promoting active fraction after purification of *E. coli* and *Salmonella* cell extracts.** The active fractions analyzed were obtained from the CM cation step of purification. See Fig. S5 and Methods. 19 proteins were identified for *E. coli* and 21 proteins for *Salmonella*; of these, 5 proteins were common.

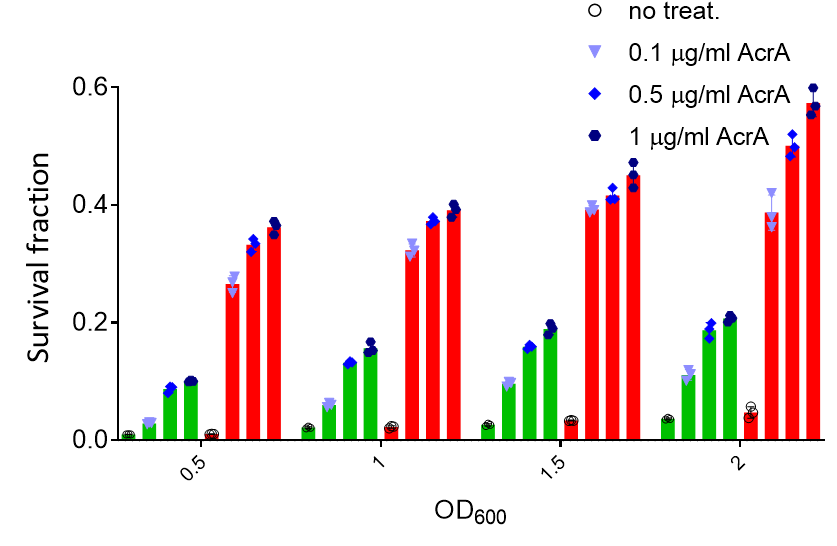

**Figure S7** | **Necrosignaling activity of purified AcrA on planktonic and swarm cells of *E. coli*.** Mid-log phase planktonic cells (green) and swarm cells (red) at various cell densities (OD_600_, x axis), either concentrated or diluted to achieve desired OD_600_, were treated with Kan^37.5^ (MIC_99_S) and Kan^20^ (~MIC_99_P) respectively for 1 h at 37^o^C with or without the addition of purified AcrA at indicated concentrations. CFU counts of survivors were used to calculate the survival fractions**.** Each of the three individual replicates are shown on each bar. The data were analyzed using a mixed model ANOVA; cell type, i.e. planktonic and swarm cells, were taken as the ‘between-subjects’ factor and a Giesser-Greenhouse correction was applied to it; cell density and treatment with AcrA were considered as repeated measures factors. The multiple comparisons were corrected using Dunnett testing keeping a two-tailed significance level of 0.05. The obtained p value of interaction was significant (<0.0001). The maximum increase in survival observed were: i) at 0.5 OD_600_, ~9% in planktonic cells compared to ~25% in swarm cells and ii) At 2.0 OD_600_, ~17% in planktonic cells compared to ~50% in swarm cells.

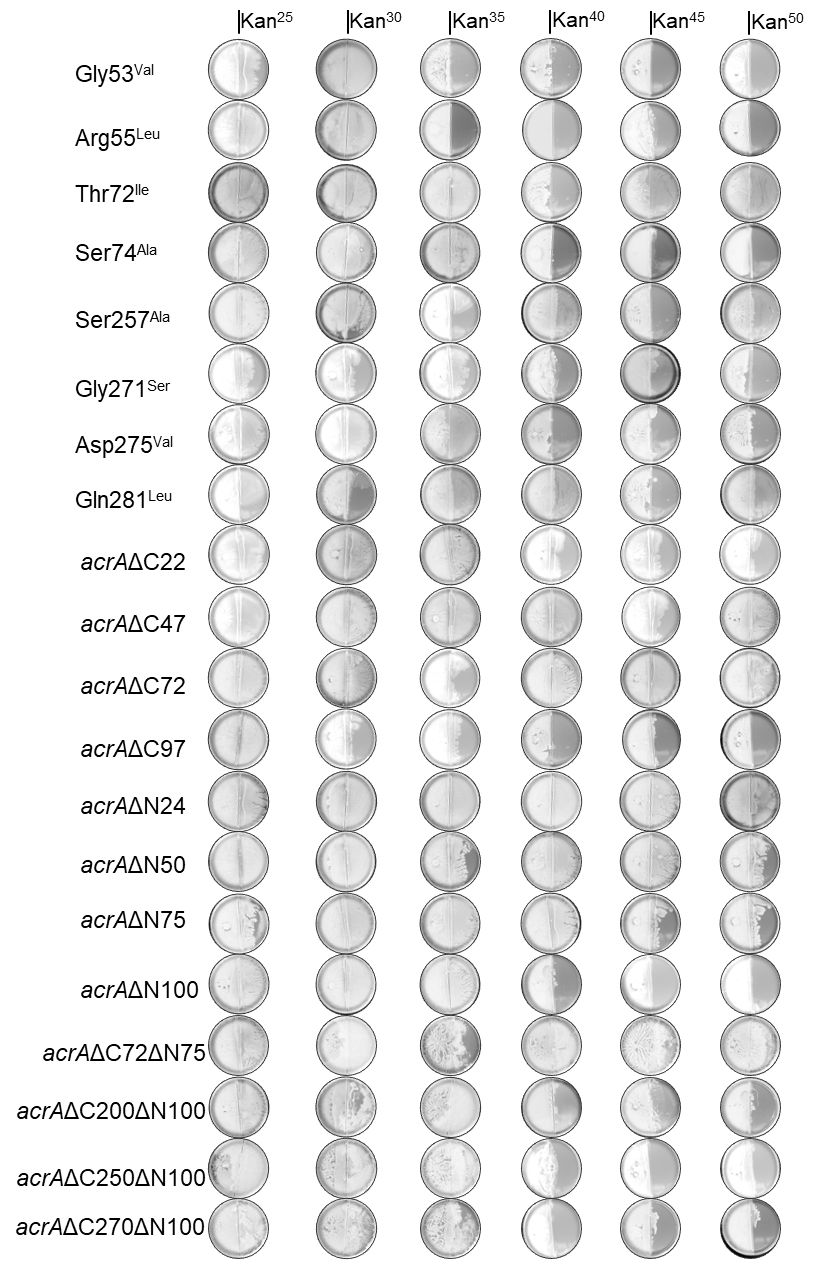

**Figure S8 | Primary data for results summarized in Fig 2c.** Indicated mutants are inoculated in the left chamber. Kan concentrations are for the right chamber.

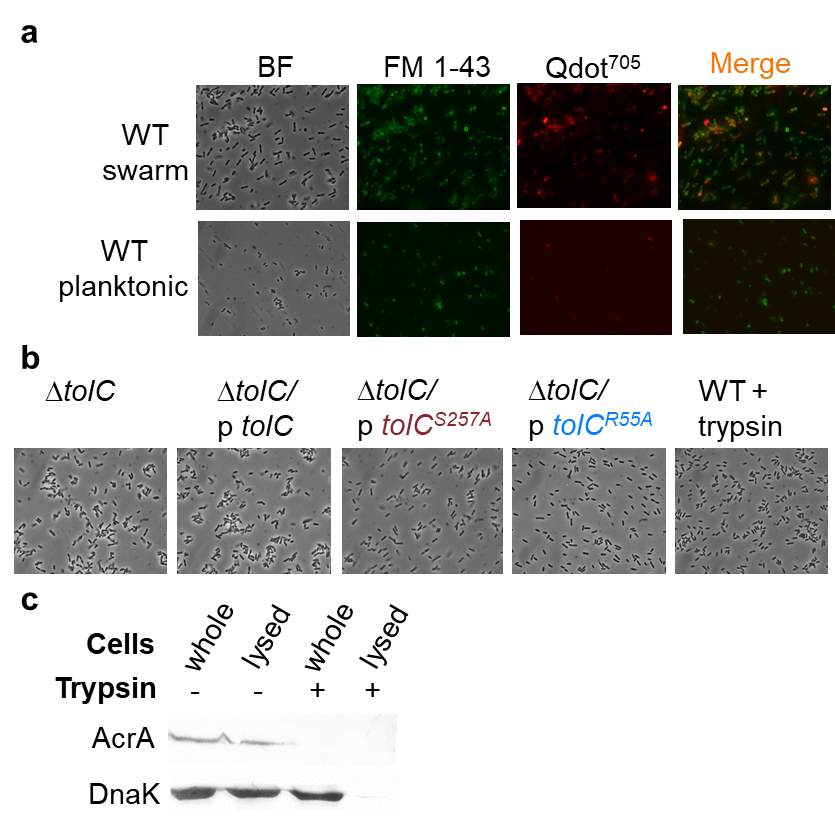

**Figure S9** | **Binding of extracellular AcrA to the outer membrane protein TolC of planktonic and swarm cells of *E. coli***. **a.** Imaging of cells probed with QDot^705^ anti-FLAG antibody and FM-143 (see Methods). The entire field of view (for each channel under the microscope) of the images that were enlarged and shown in Fig. 2d. **b**. Bright field images of swarm cells shown in Fig. 2e. Blue, SR+; maroon, SR-. **c.** Western blot analysis for detecting AcrA localization (see Methods) after trypsin digestion of cell samples used for microscopy in Fig. 2d. DnaK was probed as a cytoplasmic control. Absence of bands for AcrA (FLAG-tagged) from whole cells treated with Trypsin indicate extracellular localization of AcrA in the binding experiment in A.

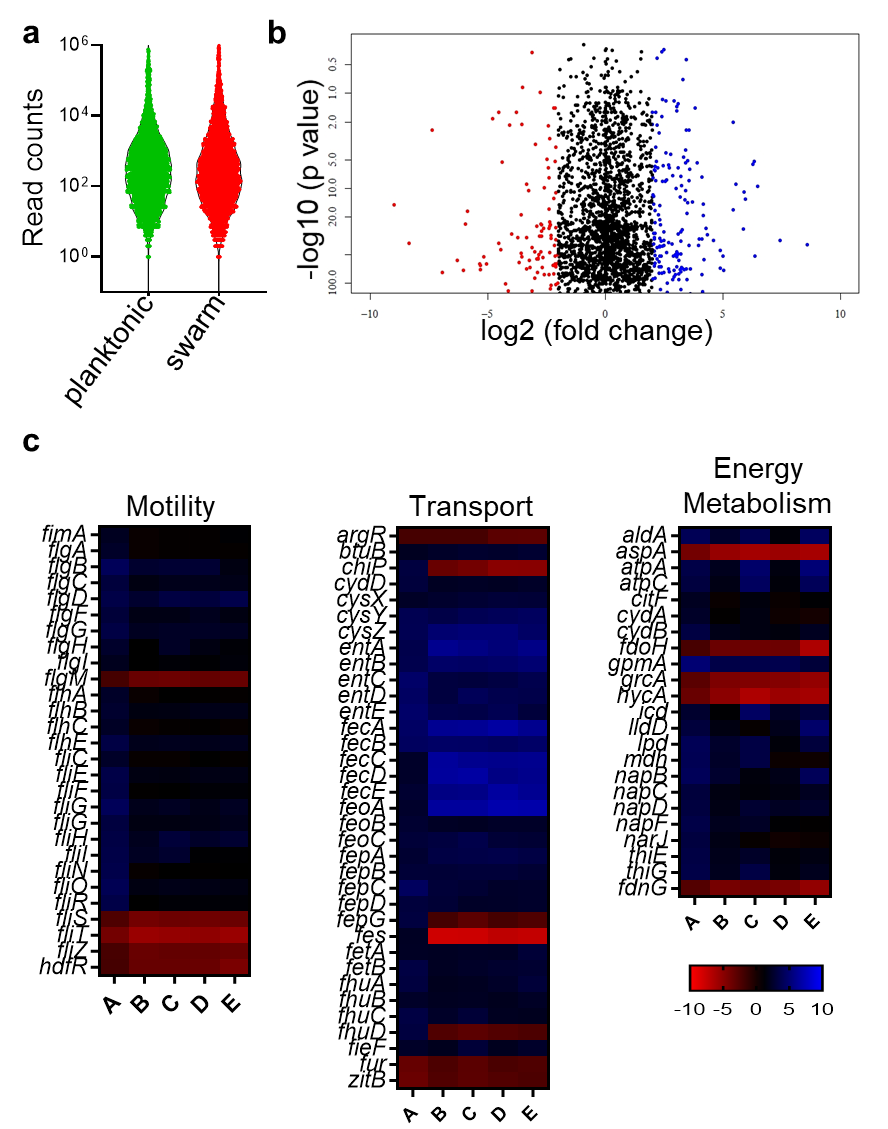

**Figure S10 |** **RNA seq analysis**. **a.** Distribution of mapped read counts for genes showing sequencing consistency across samples. **b.** A representative scatter plot showing fold change and adjusted p-value distributions of genes for swarm vs planktonic. A ±2 log2 fold change cutoff was used to identify up or downregulated genes. The blue and red dots represent log2 fold change with a value of >= 2 and <=-2, respectively. **c.** Heat maps showing changes in expression of the different gene classes (motility, transport, energy metabolism) across the following comparisons: A) Swarm vs Planktonic, B) Swarm vs Swarm+ Kan^20^, C) Swarm vs Swarm+Kan^20^+AcrA (0.1 μg/ml), D) Swarm vs Swarm+Cip^2.5^, E) Swarm vs Swarm+ Cip^2.5^+AcrA (0.1 μg/ml).

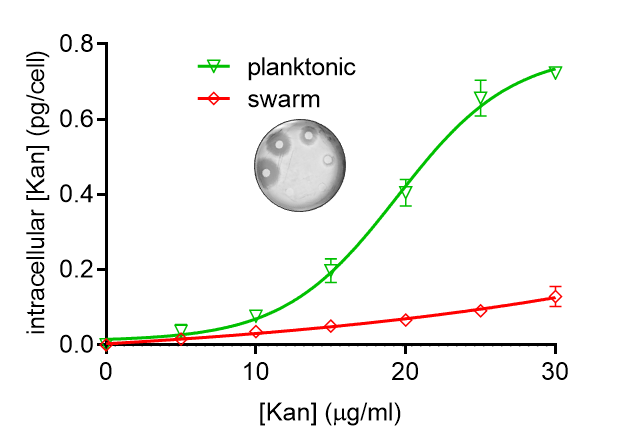

**Figure S11 |** **Disc diffusion assay to measure intracellular antibiotic concentration**. Swarm cells were collected from the right chamber of a border-crossing plate containing the indicated Kan concentrations, 2 hours after the cells crossed the border, and planktonic cultures were incubated for 2 hours in the same Kan concentrations. Cell extracts were prepared as described in Methods. Extracts were spotted on filter discs, dried for 2 hours, and placed on LB plates. Zone of inhibition was measured of these discs (for *E. coli*) extrapolated from a standard curve generated with zones of inhibition exhibited by known concentrations of kanamycin. The CFUs were counted from the cultures used for cell extract preparation. The following formula was applied to calculate intracellular [Kan]:

$\left( measured \left[ kan \right] estimated from diameter of disc \right)/(CFU of culture)$.

Swarm cells show ~ 8 fold decrease in intracellular [Kan] at Kan^30^ compared to planktonic cells.

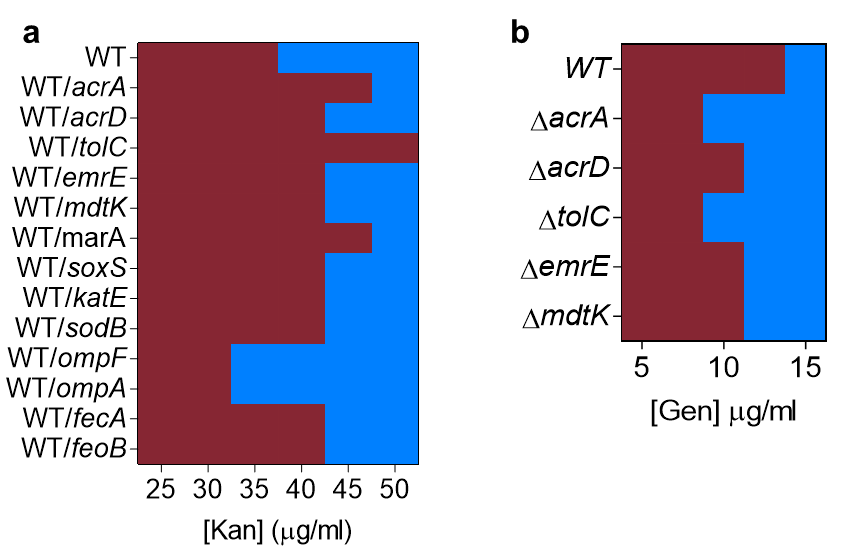

**Figure S12 |** **Genetic analysis to validate RNA seq results**. Summary of border crossing assays. Blue, SR+; Maroon, SR-. **a**. WT *E. coli* carrying plasmids from ASKA library for overexpression of selected components from different classes of efflux pumps (*acrA, acrD, tolC, emrE, mdtk*) and their regulators (*marA* and *soxS*), ROS catabolism genes (*katE* and *sodB*), and iron transport genes (*fepA and feoB*), all of which showed an increase in SR. Porins *ompF* and *ompA* reduced SR. Gen was used in the right chamber as these strains harbor a Kan^R^ marker. **b.** Deletion strains of selected efflux pump genes (*acrA*, *acrD*, *tolC*, *mdtK*, and *emrE*), inoculated on the left, reduced SR.

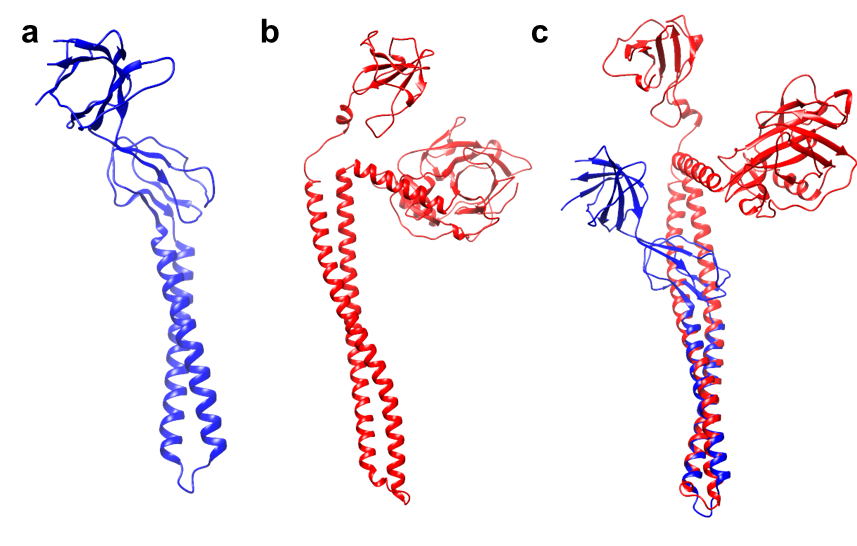

**Figure S13. Structural superposition** **of AcrA and Colicin E3.** **a**. AcrA (PDB ID: 5NG5). **B**. Colicin E3 (PDB ID: 2B5U). **C**. Superposition of A and B using Chimera matchmaker^2^. The RMSD value was 0.85 Å.

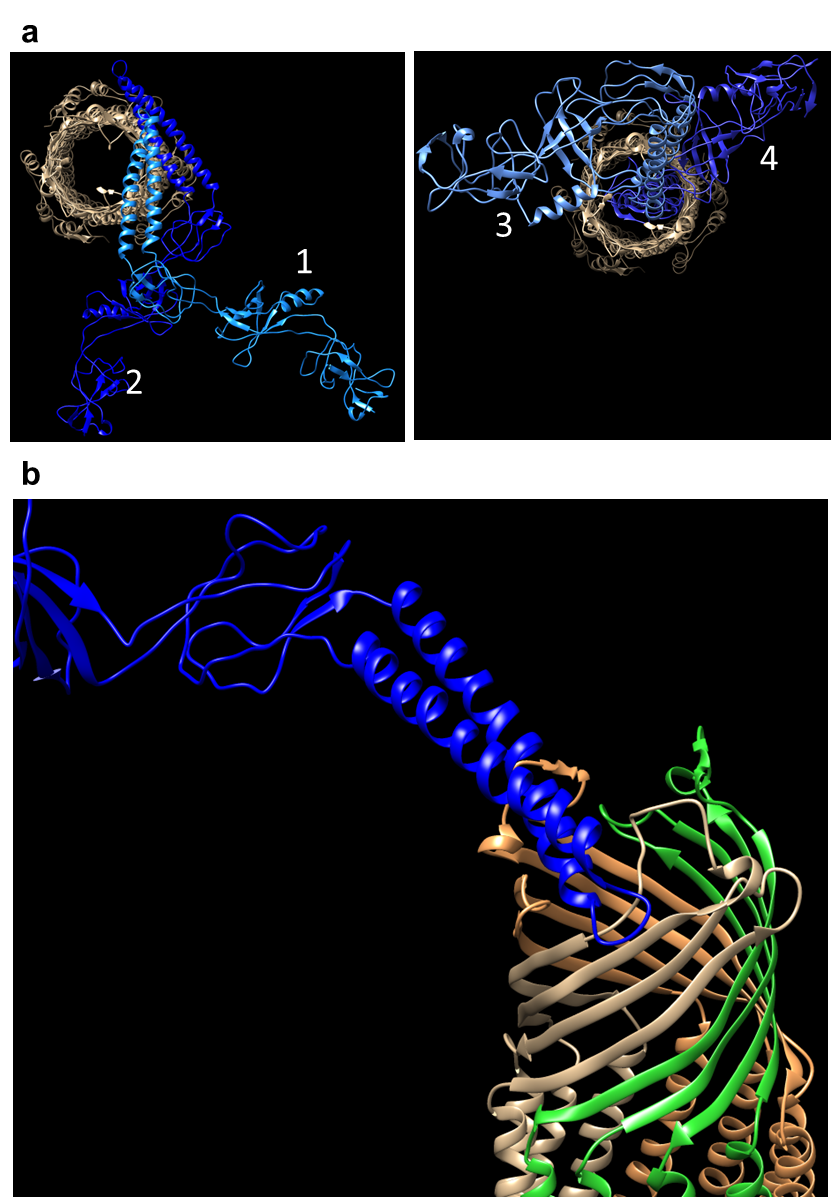

**Figure S14 | Model for AcrA-TolC binding.** AcrA and TolC structures (PDB ID 5NG5) were used to predict possible modes of their interaction (see methods). **a.** The best models are presented in decreasing order (1-4) of probability. AcrA, all shades of Blue; TolC, grey, brown, and green. **b.** Sideview of the best model (#1). It is physically not very probable for a full-length AcrA (~49 Å diameter) to enter the cell through the TolC pump (~ 29 Å diameter).

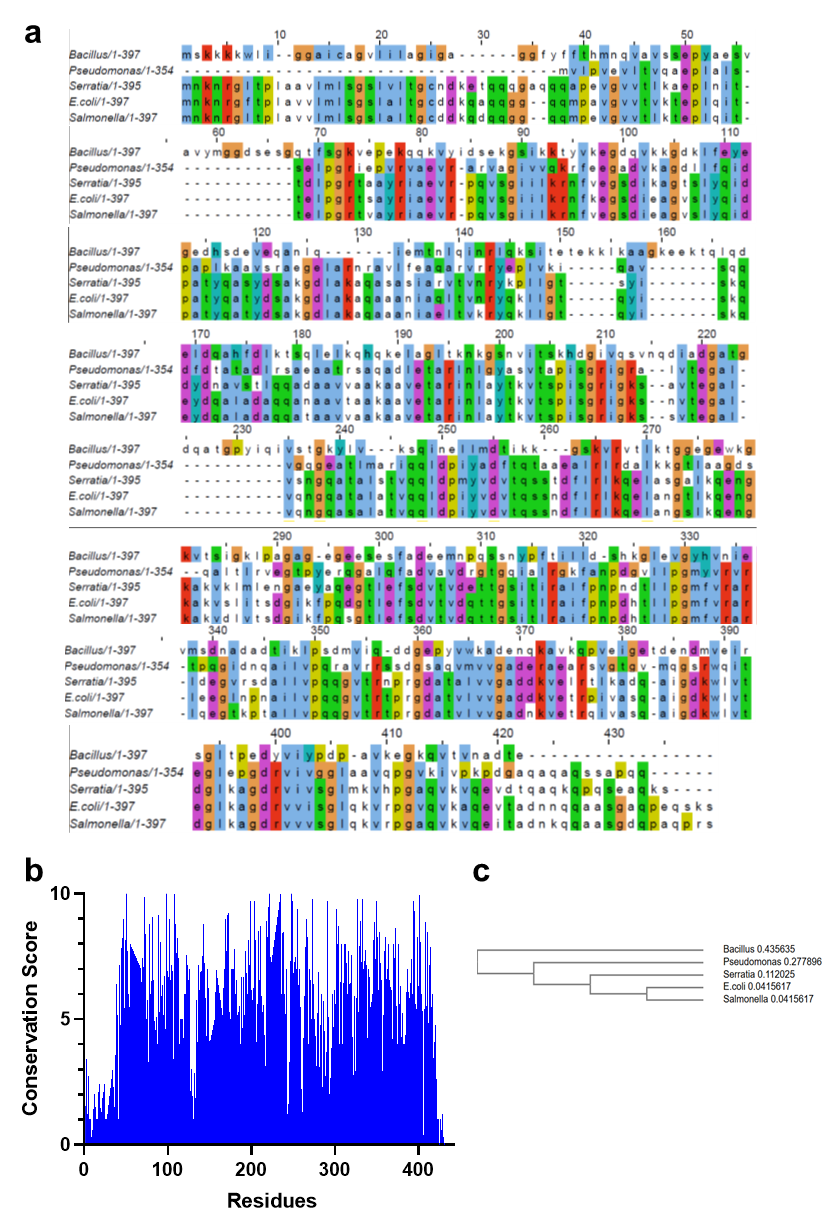

**Figure S15 | Comparison of AcrA sequences in five organisms used in this study.** **a.** Multiple Sequence Alignment (MSA) of AcrA sequences (see methods).; **b.** Graph showing variations in conservation scores across residues; numbering relates to *E. coli* AcrA sequence. **c.** A cladogram of AcrA sequences.

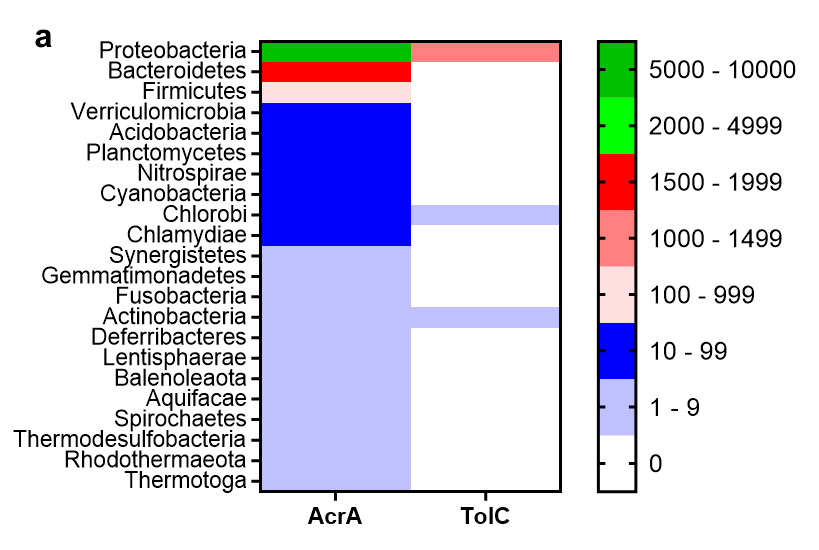

**Figure S16 | Distribution of AcrA and TolC across bacterial domain.** The sequences of AcrA and TolC of *E. coli* were used to perform a search using HMMER algorithm^3^ within bacterial taxa. The significant ‘e value’ cutoff was 10^-20^. A decrease or increase in the cutoff (10^-10^ to 10^-3^) did not alter the pattern of results significantly. The values show the number of hits obtained across different taxa.

**SI Text**

**Modeling Bacterial Response to Antibiotics**

The percent survival of the bacterial population for a given antibiotic concentration as a function of time was modeled as a set of ordinary differential equations. Initially, rate of change in a specific bacterial population was modeled as simple first-order reaction given by Eqs 1-3

$$A\underset{\to}{k_{a}}D$$

Eq. S1

$$\frac{dA_{n}}{dt}= -k_{a}\left[ A \right]_{n}$$

Eq. S2

$${[A]}_{n+1}={[A]}_{n}-k_{a}\left[ A \right]_{n}$$

Eq. S3

The population live cells, A, will convert into dead cells, D, with some rate of change, $k_{A}$. Here we have simplified all the processes that lead to changes in the bacterial population to a single term $k_{a}$ that ideally represents a convolution of both bacterial growth and death. Time-step updates to the population of A-cells is given by Eq3. The percent survival for a single population of cells is given by Eq. 4

$$S_{n}={[A]}_{n}/\left[ A \right]_{0}$$

Eq. S4

The percent survival of cells, S_n,_ is given by dividing the remaining cells at any time-point n, ${[A]}_{n}$, by the initial starting population, $\left[ A \right]_{0}$. A Gibbs sampler was built to search for suitable parameters for the rate of change for the population based on reducing the mean-square error (MSE) between the simulation and experiment over 100000 iterative steps. Simulations for a homogenous population exhibited deviation between experiment and simulation results as measured by MSE for swarming cells (10^-3^-10^-2^) and significantly better results for planktonic cells (10^-6^-10^-3^) (Fig 1a).

The failure of the simulation to reproduce the experiment was hypothesized to be the result of the simplicity of the starting assumption that there is one population of cells with a singular response rate to the antibiotics. A more complex approach was developed using a heterogeneous starting population composed of two sub-populations (A and B) that follow the same first-order kinetics shown in Eqs 1-3 but are allowed to differ in rates of change and relative starting population to one another. The survival percentages were calculated using Eq 5.

$$S_{n}=\frac{[{A]}_{n}+ {[B]}_{n}}{{[A]}_{0}+ {[B]}_{0}}$$

Eq. S5

Using the Gibbs sampler to explore the independent rates for both sub-populations, the simulation was much better able to replicate the experimental results for swarming cells with an MSE range of 10^-7^-10^-5^ _­_typically conferring a hundred to ten-thousand fold better fit over a homogenous simulation (Fig 1a). The differences between the populations usually converged into a solution where cells of one population were 10 to 100 times faster at dying, and are referred to as the fast-dying (FD) cells. Typically, the slow-dying cells (SD) where twice as numerous in starting populations than the FD cells (50-70%). Additionally, magnitude of the difference of rates of death, $\left| k_{A}-k_{b} \right|$, between the FD and SD cells decreased as a function of antibiotic concentration. In contrast, simulations on heterogeneous swimming cells showed a drastically lower rate of improvement compared to homogenous cells with MSE ranges ranging 10^-7^-10^-3^, typically indicating a 2 to 100 fold improvement in fit. The differences in MSE scores between the homogeneous and heterogeneous models are statistically significant for both swarming (p-value: 6.5 x 10^-4^) and planktonic cells (p-value: 4.5 x 10^-3^). Taken as aggregate, there is sufficient evidence that two populations of bacterial cells exist within the swarming populations that differ in initial concentrations and susceptibility to kanamycin.

**Gibbs Sampler:**

Bacterial models where built using equations S3 and S5 while determining the percent survival at discrete time steps for comparison to experimental data and MSE calculation. Each model had 2*N variables that could be altered to determine a better fit, where N is the number of sub-populations. The variables for each sub-population are the initial size of the population (A_0_ B_0_ C_0_, etc) referred to as *starting_size* and the rate of change (k_a_, k­_b_, k_c_, etc) referred to as *death_rate.* The Gibbs sampler was initiated with equal *starting_size* and random *death_rates* chosen from a uniform distribution between 0.1 and 0.001 for each sub-population. A resulting initial MSE for the model was calculated and referred to as old_MSE. To converge on a reasonable value for each variable and reduce the MSE, the following Gibbs-sampling algorithm was employed as follows for 100000 iterations:

1) Choose a variable randomly. This variable is referred to as old_variable

2) The chosen variable selects a proposed value for the variable from a normal distribution centered on the current value of the variable with a standard deviation one-tenth the size of the current value. This value will be referred to as new_variable

3) Estimate the new_MSE using the new_variable instead of the old_variable. Unselected variables will remain the same

4) If the new MSE is lower than the old MSE:
set old_variable = new_variable

Set old_MSE = new_MSE

Variables were chosen at random for each iteration, so as not to introduce an order selection bias to determining variable selections.

1	Butler, M. T., Wang, Q. & Harshey, R. M. Cell density and mobility protect swarming bacteria against antibiotics. *Proceedings of the National Academy of Sciences of the United States of America* **107**, 3776-3781, doi:10.1073/pnas.0910934107 (2010).

2	Pettersen, E. F. *et al.* UCSF Chimera--a visualization system for exploratory research and analysis. *Journal of computational chemistry* **25**, 1605-1612, doi:10.1002/jcc.20084 (2004).

3	Potter, S. C. *et al.* HMMER web server: 2018 update. *Nucleic acids research* **46**, W200-W204, doi:10.1093/nar/gky448 (2018).
